# Investigation of a Novel Mouse Model of Prader-Willi Syndrome with Invalidation of *Necdin* and *Magel2*

**DOI:** 10.1101/2024.07.24.604909

**Authors:** Pierre-Yves Barelle, Alicia Sicardi, Fabienne Schaller, Julie Buron, Denis Becquet, Felix Omnes, Françoise Watrin, Catarina Santos, Clément Menuet, Anne-Marie François-Bellan, Emilie Caron, Jessica Klucznik, Vincent Prevot, Sebastien G Bouret, Françoise Muscatelli

**Author notes:** Co-corresponding and co-lead authors, contact information: Sebastien G. Bouret and Françoise Muscatelli. Authorship note: P.-Y.B, A.S., and F.S. contributed equally to this work.

## Abstract

Prader-Willi syndrome (PWS) is a multigenic disorder caused by the loss of seven contiguous paternally expressed genes. Mouse models with inactivation of all PWS genes are lethal. Knockout (KO) mouse models for each candidate gene were generated, but they lack the functional interactions between PWS genes. Here, we revealed an interplay between *Necdin* and *Magel2* “PWS” genes and generated a novel mouse model (named “*Madin*”) with a deletion including both genes. A subset of *Madin* KO mice showed neonatal lethality. Behaviorally, surviving mutant mice exhibited sensory delays during infancy and alterations in social exploration at adulthood. *Madin* KO mice had a lower body weight before weaning, persisting after weaning in males only, with reduced fat mass and improved glucose tolerance. Delayed sexual maturation and altered timing of puberty onset were observed in mutant mice. Adult *Madin* KO mice displayed increased ventilation and a persistent increase in apneas following a hypercapnic challenge. Transcriptomics analyses revealed a dysregulation of key circadian genes and alterations of genes associated with axonal function that were also found in the hypothalamus of patients with PWS. At neuroanatomical levels, we report an impaired maturation of oxytocin neurons and a disrupted development of melanocortin circuits. Together, these data indicate that the *Madin* KO mouse is a reliable and more genetically relevant model for the study of PWS.

## Introduction

Prader-Willi syndrome (PWS) arises from the loss of expression of seven contiguous paternally inherited genes located in the 15q11-q13 region, including the *MAGEL2* and *NECDIN* genes. All the PWS genes are regulated by genomic imprinting, which is an epigenetic process where only one allele is expressed in a parent-of-origin dependent manner. PWS is a complex neurodevelopmental disorder characterized by a lifelong spectrum of phenotypic features starting with severe feeding difficulties and respiratory distress at birth, early sensory deficits and hypotonia, followed by growth deficiency, hypogonadism and delayed puberty, short stature, excessive weight gain with severe hyperphagia, and cognitive and behavioral problems throughout life (1, 2). Despite extensive clinical trials, there are no effective treatments for PWS, and comprehensive pathophysiological mechanisms have not yet been clearly identified, although converging evidence suggests that the PWS phenotype might result from hypothalamic dysfunctions (2, 3). Importantly, point mutations in the paternal allele of *MAGEL2* only are responsible for the Schaaf-Yang syndrome (SYS) that has a phenotypic overlap with PWS but with a more severe autistic phenotype in adolescence and adulthood (4, 5). Altogether, these results suggest that the loss of function of *MAGEL2* in SYS shares common overlapping symptoms with PWS.

Animal models for PWS are instrumental in understanding mechanisms and identifying novel pathways involved in the pathophysiology of PWS. The mouse chromosome 7C presents a conserved synteny to the human PWS region. However, mouse models with inactivation of all PWS genes display a 100% lethality rate within the first week after birth and have, therefore, not been very useful for understanding postnatal symptoms (6, 7). Mouse knockout (KO) models for single candidate genes have also been generated (8). However, these single KO models have limitations as it is likely that the PWS phenotype is the result of the lack of expression of several genes that are co-expressed in the same brain regions (9) and that may interact with each other, creating a more complex and integrative phenotype.

Among the genes inactivated in PWS, *NECDIN* and *MAGEL2* are of particular interest. *Necdin* and *Magel2* single KO mouse models display several distinct phenotypes mimicking part of the PWS clinical features, although there is variability among the different models depending on the genomic construction. *Magel2* KO mice exhibit suckling deficits at birth (10), growth retardation (10), altered metabolism (11, 12), circadian activity disturbances (13), deficits in cognitive, social, and parental behaviors (14–17). Our teams and others also reported impaired hypothalamic regulation in *Magel2* KO mice with abnormal oxytocin (OT) maturation (10, 14, 18, 19), and disrupted development and function of pro-opiomelanocortin (POMC) neurons (19, 20). *Necdin* KO mice display variable lethality after birth (21, 22) due to respiratory distress (23), growth retardation, motor deficit in infancy (24), sensory deficits (25), high scraping, cognitive alterations (22) and alterations of social and circadian behaviors (26, 27). At the neuroanatomical level, *Necdin* KO mice display a reduction in the number of OT and gonadotropin-releasing hormone (GnRH)-producing neurons, alterations in perinatal serotonergic metabolism and development (22, 28), and alterations in clock gene expression (26).

*MAGEL2* and *NECDIN* belong to the MAGE (Melanoma Antigen Gene Expression) gene family (29). They are physically close in the genome (30kb), intron-less, and have probably evolved through sequential retrotransposition events (9). In addition, they are co-expressed in many brain structures, including in the developing hypothalamus (9, 30, 31). At the molecular level, both proteins act through a ubiquitin-dependent mechanism to turn over and recycle proteins (32). Overall, their molecular function and expression pattern suggest that their roles in cellular processes may partially overlap (32). Therefore, an animal model with the combined loss of *Magel2* and *Necdin versus* a single invalidation of each gene should reveal the complex interaction of these two genes and avoid the potential functional redundancy or compensatory mechanism between these two genes. This model would therefore be more relevant to study the complexity of PWS.

In the present study, we investigated the interplay between *Necdin* and *Magel2* genes, generated a novel mouse model with a deletion including both *Magel2* and *Necdin* genes, and provided a comprehensive characterization of the behavioral, physiological, neurodevelopmental, and transcriptomic alterations of this model.

## Results

### Co-expression and co-regulation of *Magel2* and *Necdin* genes and generation of a mouse model with a deletion including both genes

We previously found that *Necdin* and *Magel2* mRNAs were highly expressed in the developing brain (9). Here, we showed a striking overlapping expression pattern of both genes in the embryonic and adult brain (Figure 1A-B). Our previous studies (9) and genomic analysis (https://genome.ucsc.edu) revealed that both genes share a common enhancer (Figure 1C). We have previously generated two mouse models in which the promoter and 5’ of the coding region of *Necdin* (*Ndn^tm1-Mus^*) (22) or *Magel2* (*Magel2^tm1-Mus^*) (10) were deleted preventing the expression of *Necdin* and *Magel2* transcripts, respectively. In the present study, we used RT-qPCR and found that *Necdin* mRNA was over-expressed in *Magel2* KO brains, and *Magel2* mRNA was over-expressed in *Necdin* KO brains (Figure 1D). We also showed by *in situ* hybridization that *Magel2* transcript was overexpressed in *Necdin* KO brain (Figure 1E) and Necdin immunoreactivity was increased in the brains of *Magel2* KO mice (Figure 1F). Together, these data show that *Magel2* and *Necdin* are spatio-temporally co-regulated, and share a putative enhancer, with functions that partially overlap.

**Figure 1.**
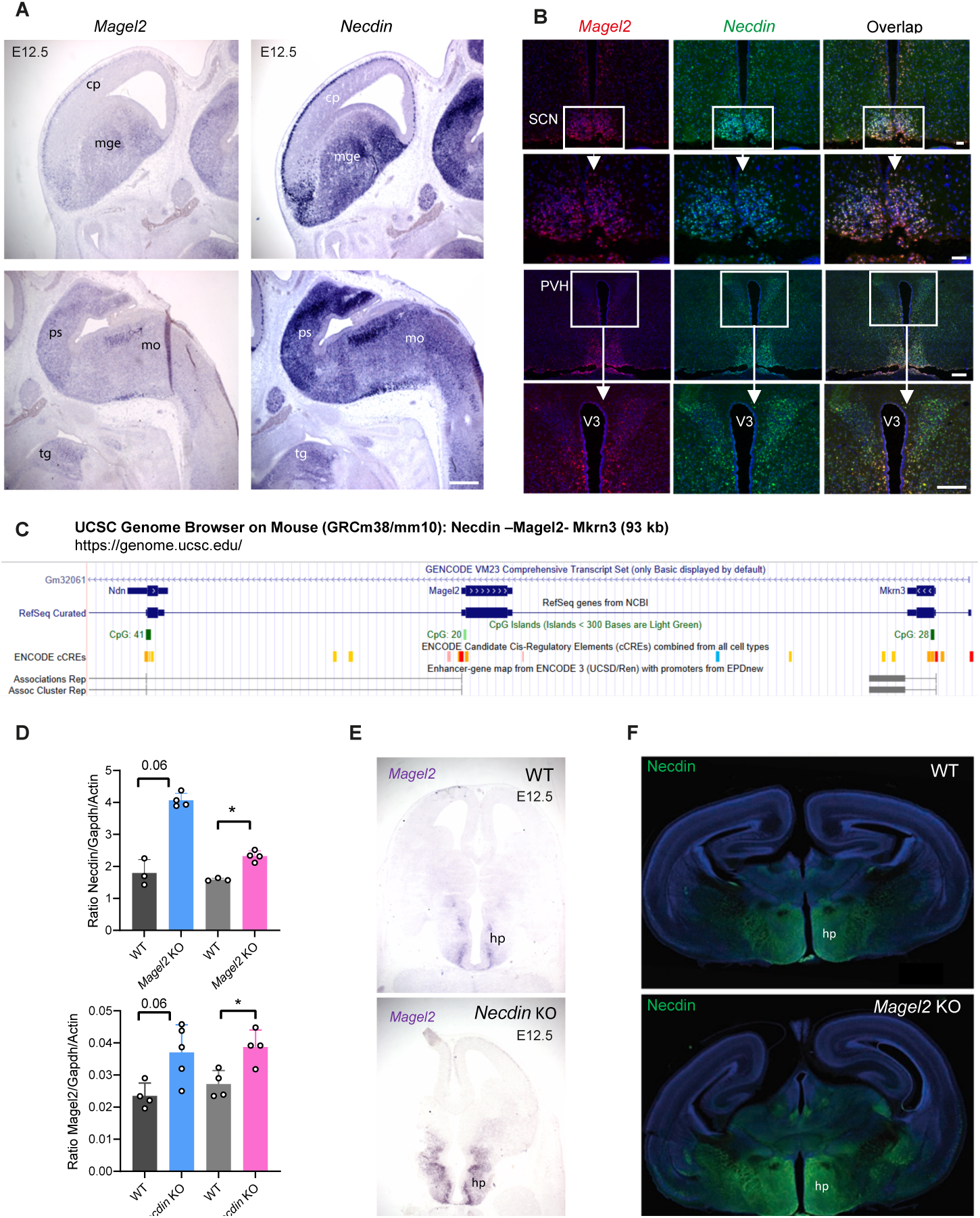
Co-expression and co-regulation of *Necdin* and *Magel2* genes in the mouse brain. (**A,B**) Images showing *Necdin* and *Magel2* mRNA-expressing cells in (**A**) the forebrain and brainstem of E12.5 mice and (**B**) in the hypothalamus of adult mice. (**C**) Map of the 93kb mouse genomic region including *Necdin, Magel2* and *Mkrn3* genes, ENCODE Cis regulatory elements, and associations between enhancers and promoters of genes, extracted from UCSC Genome Browser. A co-regulation of *Magel2* and *Necdin* via shared enhancer is proposed. (**D**) RT-qPCR analysis showing relative levels of *Necdin* and *Magel2* mRNA in wild-type (WT), *Magel2* KO, or *Necdin* KO male and female mice at P0 (n = 3-5 animals per group). (**E**) Images showing *Magel2* mRNA-expression on horizontal brain sections (at the level of the presumptive hypothalamus) of WT and *Necdin* KO embryos at E12.5. (**F**) Images showing Necdin immunoreactivity in coronal sections at the level of the hypothalamus of WT and *Magel2* KO mice at P0. Data are presented as mean ± SEM. Statistical significance between groups was determined by a Mann-Whitney test (D). \**P* < 0.05. Scale bars: 50 μm (A,E), 20 μm (B), 500 μm (F). cp, cortical plate; hp, hypothalamus; mge, median ganglion eminence; mo, medulla oblongata; ps, pons; tg, tongue; PVH, paraventricular nucleus; SCN, suprachiasmatic nucleus; V3, third ventricle.

Based on the findings described above, it appears that the combined loss of *Magel2* and *Necdin,* by reflecting more accurately the genetics of PWS, will be a more appropriate model to study PWS-like phenotype. We, therefore, generated a large deletion including both *Necdin and Magel2 genes* (hereafter called “*Madin*” KO mice for the contraction between *Magel2* and *Necdin*) using both *Ndn^tm1-Mus^* and *Magel2 ^tm1-Mus^* single KO mice and an *in vivo* chromosomal rearrangement based on the Cre-loxP system (Figure 2A). This strategy allowed a somatic or meiotic trans-allelic recombination between loxP sites in a transgenic mouse expressing the *Hprt*-Cre recombinase (33). We used it to create a recombination between both loxP sites located in the same direction but associated with the mutated *Necdin* allele or the mutated *Magel2* KO allele, both alleles being on a different chromosome 7. We created a novel allele with a deletion that includes *Necdin*, the 30 kb-fragment between *Necdin* and *Magel2*, the *Magel2* promoter, and half of the *Magel2* coding part (Figure 2A). Notably, the 32 kb deleted fragment contained no gene, miRNA, or identified regulatory elements except the enhancers and promoters associated with the *Necdin* and *Magel2* genes. We screened and validated the recombined *Madin* allele by PCR using *Necdin* and *Magel2* primers, and we sequenced the recombined allele confirming a recombination at the loxP sites and the expected deletion (Figure 2B-C). Because *Magel2* and *Necdin* are regulated by genomic imprinting resulting in transcriptional silencing of the maternal allele and expression of the paternal allele only, all studies described below were performed on heterozygous mice with a mutated paternal allele and a wild-type yet silent maternal allele (+m/-p), considered as KO. We measured *Magel2* and *Necdin* mRNA levels in the *Madin* KO heterozygous mice (+m/-p) using RT-qPCR and confirmed the loss of expression of both transcripts (data not shown). We also confirmed the loss of Necdin protein expression in *Madin* KO brains using immunohistochemistry (Figure 2D). The loss of Magel2 protein expression could not be checked due the lack of specific MAGEL2 antibodies.

**Figure 2.**
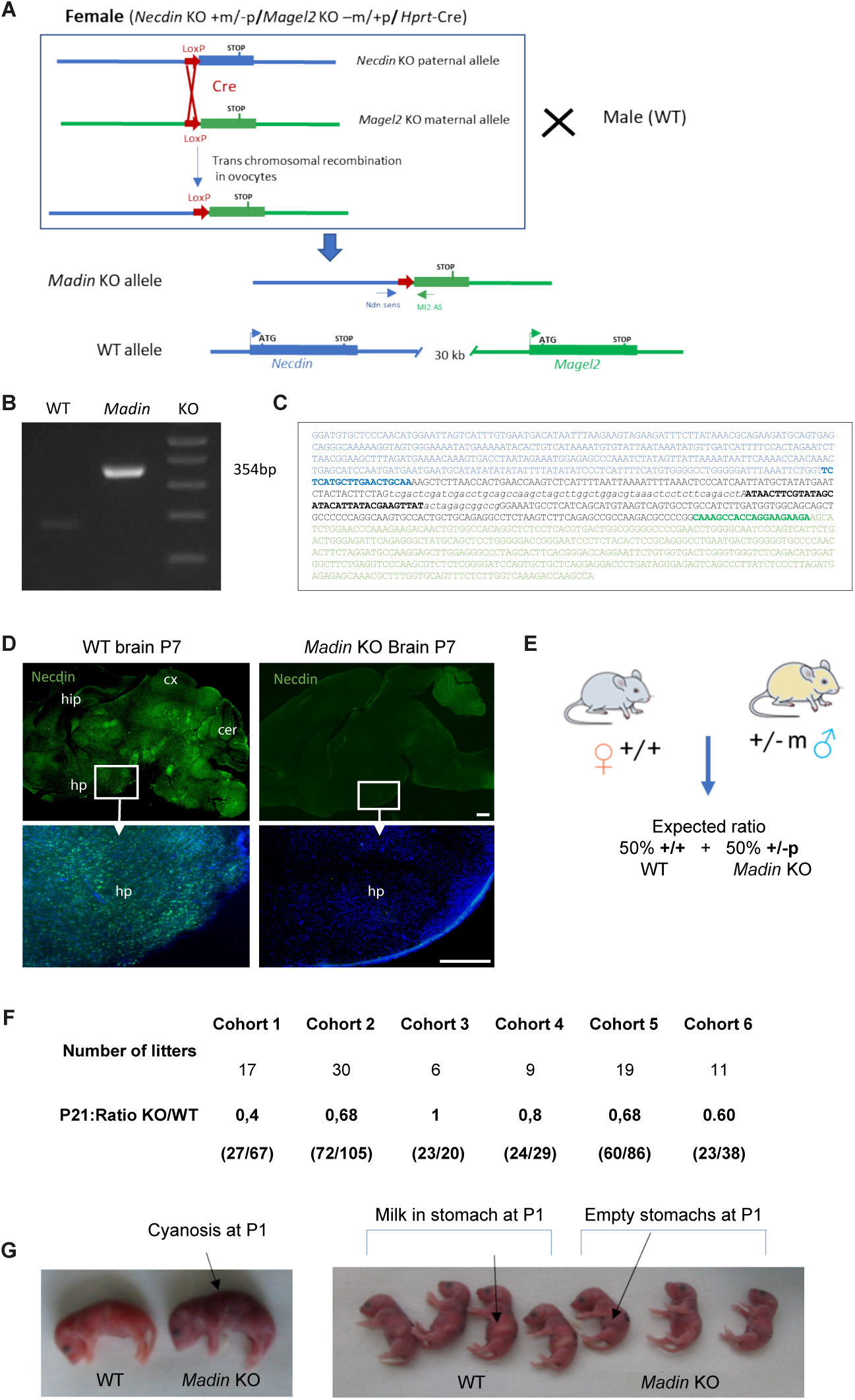
Construction and validation of the *Madin* KO mouse model. (**A**) Strategy to obtain a 32 kb deletion including both the *Necdin* and *Magel2* genes using a trans-allelic recombination approach. First, a female mouse containing one maternal allele with the *Magel2* deletion, one paternal allele with the *Necdin* deletion, and containing the *Hprt*-Cre gene was created using both *Necdin^tm1-Mus^* and *Magel2 ^tm1-Mus^* KO models and a transgenic mouse line expressing the Cre recombinase under Hprt promoter. We then crossed these female mice with wild-type (WT) male mice and screened the litter for the recombinant *Madin* KO allele using PCR with *Necdin* sense and *Magel2* antisense primers. The *Madin* allele being created in the ovocytes. (**B**) PCR product obtained from the recombinant *Madin* allele (354bp) in the F1 generation. (**C**) Sequence of the recombined *Madin* allele with the *Necdin* upstream sequence (blue) and *Ndn* sense primer (bold blue), the LoxP sequence (bold black), and *Magel2* sequence (green), and Ml2 anti-sense primer (bold green). This sequence validates the recombination at the loxP sites. (**D**) Images showing Necdin immunoreactivity on sagittal sections of WT and *Magel2* KO mouse brains at P7. (**E**) Breeding strategy to generate *Madin* KO and WT litters with an expected ratio of 1 *Madin* KO /1 WT. (**F**) Ratio of *Madin* KO and WT mice at P21 from six cohorts produced in the different laboratories performing experiments for this study. (**G**) Photos showing cyanosis and lack of milk in the stomach associated to the death of *Madin* KO pups at P1. Scale bar: 400 μm. cer, cerebellum; cx, cortex; hp, hypothalamus; hip, hippocampus.

We generated several cohorts of *Madin* heterozygous and wild-type mice in different laboratories. Although the expected Mendelian ratio of our cross was 1:1, we often observed a reduced number of *Madin* KO mice compared to wild-type animals (Figure 2E). In 5 out of 6 cohorts generated (representing 92 litters), we found a reduction of 40% (ranging from 20% to 60%) of *Madin* KO mice compared to wild-type animals (Figure 2F). The variability in the postnatal lethality observed in *Madin* KO pups appeared to be correlated with the level of sanitary status of the animal facility in which animals were housed: when animals were housed in a pathogen-free animal facility and fed sterile food, the phenotype was much less severe (with 0% lethality) than when animals were housed in a conventional animal facility (with 60% lethality in mutant mice *versus* control littermates). The lethality occurred during the first day of life. Monitoring neonates shortly after birth revealed cyanosis in *Madin* KO mice (Figure 2G), suggesting a lack of proper oxygenation due to respiratory dysfunction as observed in *Ndn^tm1-Mus^* KO mice (28, 34). *Madin* KO neonates also lacked milk in their stomachs (Figure 2G) as previously observed in the *Magel2 ^tm1-Mus^* KO mouse model (10). Together, these data indicate that the possible causes of death in *Madin* KO newborns include respiratory defects and/or feeding deficits at birth.

### *Madin* KO mice develop sensory alterations in infancy and social exploration deficits during adulthood

PWS is a neurodevelopmental syndrome and patients with PWS present symptoms at birth evolving with age. Moreover, *Necdin* and *Magel2* are highly expressed in the developing brain (Figure 1A,B). Based on these observations, we characterized the phenotype of *Madin* KO mice beginning at birth. We first used 11 reliable tests established by Roubertoux et al. (35) to assess the sensory and motor abilities during the first two weeks of postnatal life (Supplemental Figure 1A). Each test was performed at specific postnatal ages defined by the critical period during which the response is established in 100% of control mice in the mouse strain that we used (*i.e.*, C57Bl6/J) (Supplemental Figure 1B). We compared *Madin* KO to wild-type pups from the same litters. Since we did not observe differences between males and females, we pooled both sexes.

We found differences between mutant and wild-type mice in 5 out of the 11 tests. *Madin* KO pups were less efficient in righting response, rooting response, and paw position on the floor (Figure 3A-C). The righting response reveals motor and sensory (proprioception) abilities and its development was delayed in *Madin* KO pups at P4 and P8 (Figure 3A). The rooting reflex involves cranial sensory nerves (Vth); this reflex disappeared in wild-type mice across postnatal development but it was more frequent in *Madin* KO pups at P7 and at P12 (tendency) compared to wild-type mice (Figure 3B). The paw position test is mainly a sensory test (proprioception, touch) and revealed a delay in sensory development in *Madin* KO pups (Figure 3C). In contrast, mutant mice performed better in the bar-holding test at P11 (Figure 3D). This is a motor test, requiring muscle strength. Interestingly, *Madin* KO pups also performed better in the pulling up on bar test at P12 (Figure 3E), *i.e.*, they were more efficient in bringing back the bar to them and using their hind legs. However, they were less efficient in standing up on the bar at P15, requiring more sensory abilities (Figure 3E). However, mutant and wild-type pups performed similarly in the vertical climbing test, and the two other tests involving vestibular and motor activity (*i.e.*, Climbing the slope and Cliff avoidance) (Supplemental Figure 2C-E). Similarly, the age of eye opening and auditory canal opening, which normally occurs during a specific time window between P13 and P14, was similar between *Madin* KO and wild-type mice (Supplemental Figure 2 A,B). Altogether, these results indicate that *Madin* KO pups presented a delay in sensory (tactile and proprioception) maturation but not in motor abilities as they actually displayed an increased muscular strength.

**Figure 3.**
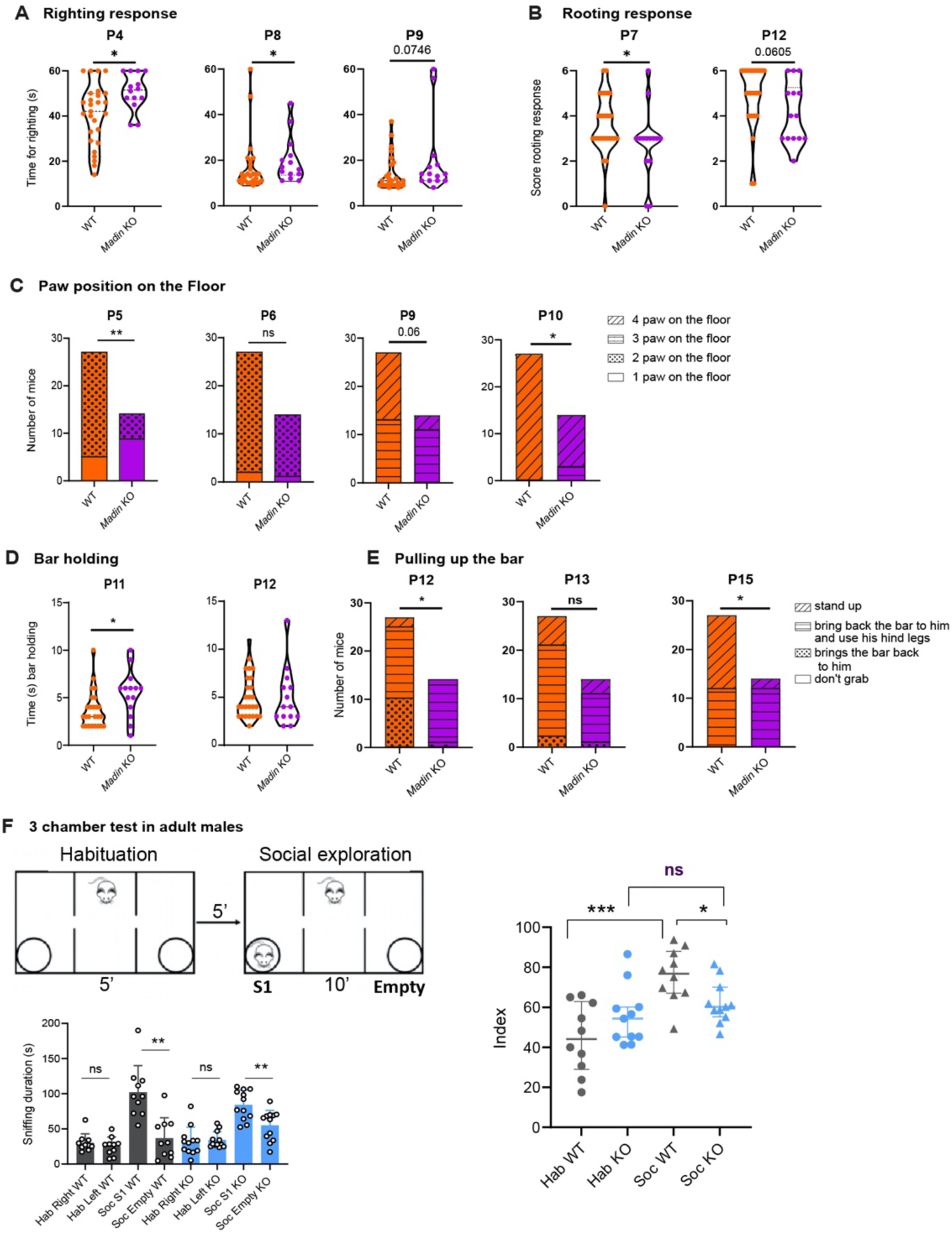
*Madin* KO mice display early sensory alterations and social exploration deficits during adulthood. (**A**) righting response in WT and *Madin* KO mice at P4, P8, and P9 (n = 14-27 animals per group), (**B**) rooting response in P7 and P12 WT and *Madin* KO pups (n = 14-27 animals per group), (**C**) paw position test in WT and *Madin* KO mice at P5, P6, P9, and P10 (n = 14-27 animals per group), (**D**) holding bar test in P11 and P12 WT and *Madin* KO pups (n = 14-27 animals per group), (**E**) test assessing the pulling up on the bar after hanging in P12, P13, and P15 WT and *Madin* KO pups (n = 14-27 animals per group). These tests evaluate sensory motor abilities. (**F**) 3-chamber test in adult WT and *Madin* KO mice (n = 14- 27 animals per group) reporting the interaction time (*i.e.*, sniffing duration in seconds) between mice measured during habituation (Hab, left or right empty grid) or in the context of social exploration (Soc, empty grid *versus* novel mouse S1). Data are presented as mean ± SEM. Statistical significance between groups was determined by a Mann-Whitney test (A,B,D), a Chi^2^ test (C,E), or a Wilcoxon matched pairs test and Mann Whitney test (F). * *P* <0.05, \*\**P* < 0.01

We performed a series of behavioral tests in adult mice that were previously done in the *Magel2* KO mouse model. No differences were observed between *Madin* KO and wild-type mice in the Rotarod test that measures motor function and coordination (Supplemental Figure 3A) and in the open field (OF) test, assessing the spontaneous locomotor activity in a novel environment (grooming, rearing, traveled distance) and giving information about anxiety-related behavior (the time spent in the center zone *versus* border of the arena) (Supplemental Figure 3B). Anxiety behavior was not affected in the mutant mice as shown with the elevated plus maze test that measured the number of entries and time spent in open arms (Supplemental Figure 3D). In addition, we did not observe differences in the spontaneous social interactions between two mice moving freely in the arena of the OF by measuring the frequency, latency to the first interaction, and duration of interaction (Supplemental Figure 3C). In the object recognition test, *Madin* KO mice spent more time exploring the novel object (sniffing time) than the familiar object, similar to wild-type mice. However, mutant mice did not travel shorter distances to reach the novel object, as expected and observed in wild-type animals (Supplemental Figure 3E). These results suggest that when close to the novel object, *Madin* KO mice showed a greater interest in the novelty than in the familiar object, but when far from the novel object, they did not preferentially move towards the novel object, suggesting a recognition deficit or interest alteration.

Sociability was evaluated, in male mice only, by testing the preference for a congener placed in a wire cage as compared to an empty wire cage (three-chamber test, social recognition task). Wild-type and *Madin* KO mice spent more time interacting with a congener compared with the empty grid (Figure 3F, sniffing duration). However, the index of sociability was significantly lower in *Madin* KO mice and is even similar to the index of habituation (Figure 3F, Index) suggesting an alteration of social recognition (and sociability) in *Madin* KO compared with wild-type mice.

### Effect of the combined deletion of *Magel2* and *Necdin* on body weight, body composition, energy and glucose homeostasis

To evaluate the physiological consequences of the loss of *Magel2* and *Necdin*, we first measured the body weight of *Madin* KO and wild-type mice throughout postnatal life. Mutant mice displayed a lower body weight, starting at P1 and continuing until P15 (Figure 4A). Notably, the difference in body weight at weaning in female *Madin* KO mice appeared to be associated with a smaller body length (Figure 4J). The differences in body weight and length in female mice were specific to the pre-weaning period, as *Madin* KO females had normal body weight curves and body length after weaning although they were leaner (Figures 4K and 4L- N). In contrast, male *Madin* KO mice displayed lower body weights but a normal body size from weaning (P21) to six months of age (P168) (Figure 4B-C and 4D). The differences in body weight in *Madin* KO males were associated with changes in body composition, as revealed by a lower fat mass and a higher lean mass compared to wild-type mice (Figure 4E-G). However, we did not find differences in fat mass in female mutant mice (Figure 4M). Also, the mean and maximal brown adipose tissue temperature was normal in male and female *Madin* KO mice (Figure 4H-I and 4P-Q).

**Figure 4.**
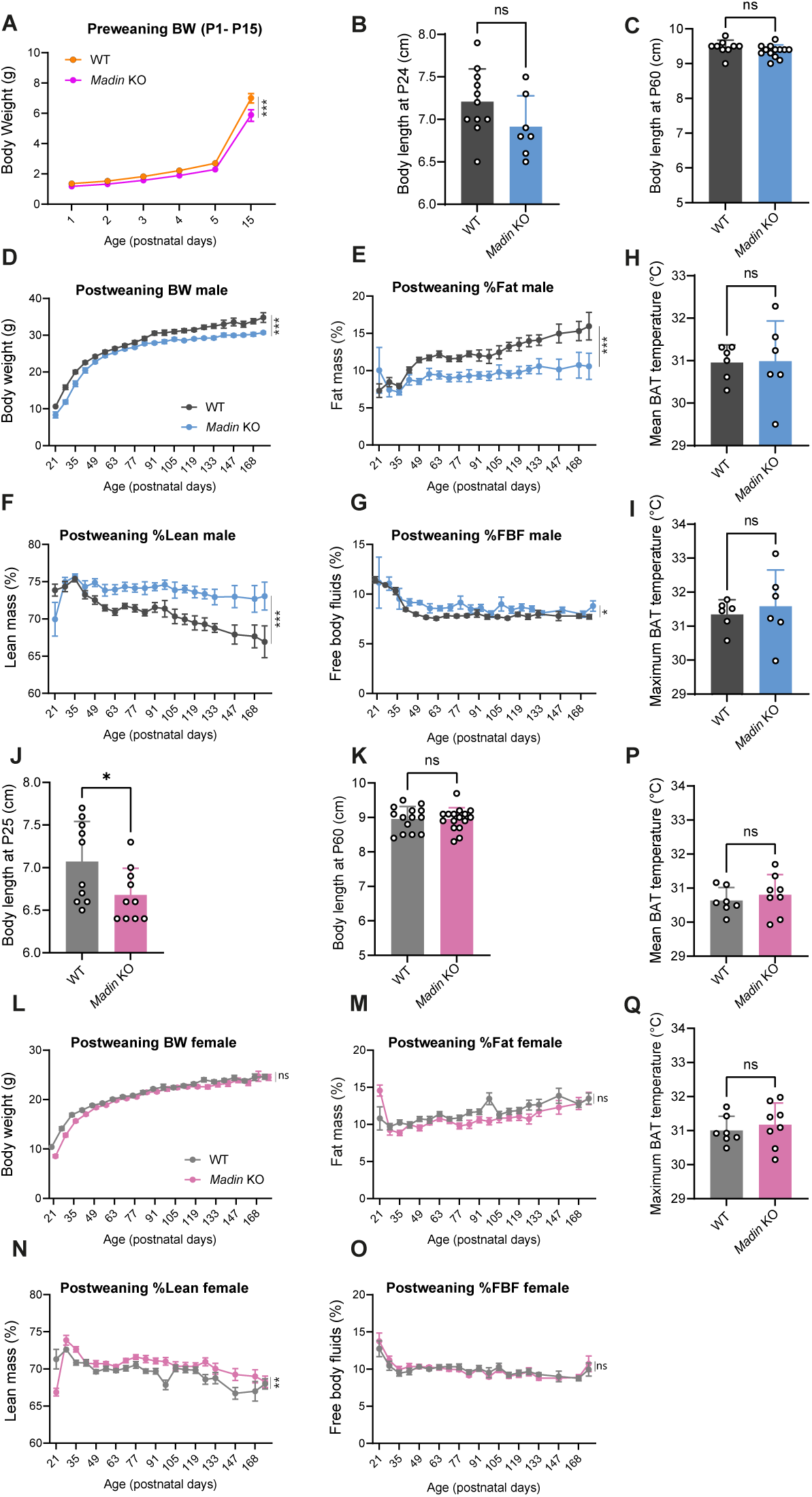
Sexually dimorphic effects of *Magel2* and *Necdin* deficiencies on growth curves and body composition. (**A**) Body weight from birth (P0) to P15 of *Madin* KO and wild-type mice (n = 13–28 animals per group). Body length at P24 and adulthood (P60) of (**B, C**) male and (**J, K**) female *Madin* KO and wild-type mice (n = 13–7 animals per group). Body weight (**D, L**), fat mass (**E, M**), lean mass (**F, N**), and free body fluids (**G, O**) of (**D-G**) male and (**L-O**) female *Madin* KO and wild-type mice from weaning (P21) to P168 (n = 13–18 animals per group). (**H, P**) Mean and (**I, Q**) maximum brown adipose tissue temperature in (**H, I**) male and (**P, Q**) female *Madin* KO and wild-type mice at P120 (n = 6 animals per group). Data are presented as mean ± SEM. Statistical significance between groups was determined by Mixed-effect test (A,D,E,F,G,L,M,N,O), Student’s t-test (B,C,J,K,L,Q,R) or a Mann-Whitney test (H,I) . \**P* ≤ 0.033., \*\**P* ≤ 0.002, \*\*\**P* ≤ 0.0002.

It is known that body composition is a factor that could influence glucose regulation and it has been reported that patients with PWS have improved glucose metabolism (36). We, therefore, measured several indices of glucose homeostasis in *Madin* KO and wild-type mice. Both male and female *Madin* KO mice exhibited lower fasting glycemia levels than their respective littermates (Figure 5A and 5G). Additionally, male, but not female, *Madin* KO mice displayed an improved glucose tolerance after a glucose challenge (Figure 5B-C and 5 H-I).

**Figure 5.**
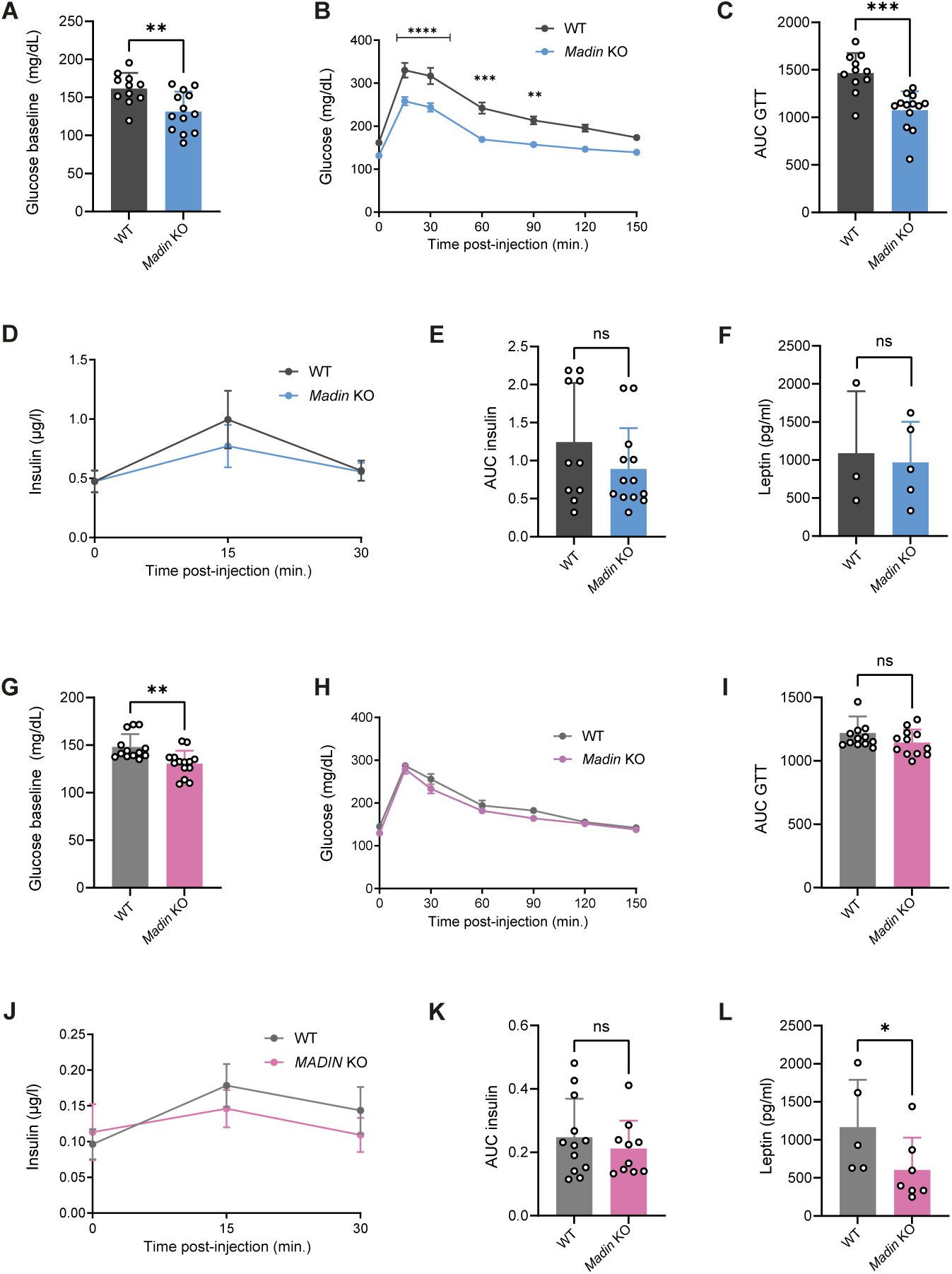
The combined loss of *Magel2* and *Necdin* alters glucose metabolism in males. (**A, G**) Basal glycemia, (**B, H**) Glucose tolerance tests (GTT), (**C, I**) areas under the GTT curve, (**D, J**) Serum insulin levels during (GTT) and (**E, K**) areas under the curve in (**A-E**) male and (**G-K**) female *Madin* KO and wild-type mice at P130 (n = 11–13 animals per group). (**F, L**) Serum Leptin concentration in (**F**) male and (**L**) female *Madin* KO and wild-type mice at P160- P240 (n = 3–4 animals per group). Data are presented as mean ± SEM. Statistical significance between groups was determined by Mann-Whitney test (A,G,E,I,K,L), or two-way ANOVA with the Geisser-Grennhouse correction following by a Tukey’s multiple comparisons test (B,H) or a Sidak’s multiple comparisons test (D,J). \**P* ≤ 0.033, \*\**P* ≤ 0.002, \*\*\**P* ≤ 0.0002., \*\*\*\**P* ≤ 0.0001

The increased glucose tolerance appeared independent of improved insulin secretion, as *Madin* KO mice had normal insulin levels during the glucose tolerance test (Figure 5D-E and 5J-K). We also conducted a comprehensive assessment of the energy balance regulation of our new mouse model. No alterations in food intake, energy expenditure, respiratory exchange ratio (RER), or xy locomotor activity were found in either male or female *Madin* KO mice compared to their wild-type littermates (Supplemental Figure 4A-H). Similarly, metabolic responses to fasting and refeeding appeared normal in mutant mice (Supplemental Figure 5). Nevertheless, the combined loss of *Magel2* and *Necdin* impacted z-rearing behavior, with a greater frequency of z-rearing during the light phase in male, but not female, *Madin* KO mice (Supplemental Figure 4I-J).

### Delayed puberty onset in mice lacking both *Magel2* and *Necdin*

Hypogonadism and delayed puberty are often reported in patients with PWS (1). We then characterized sexual maturation in *Madin* KO mice by monitoring the age of balano-preputial separation in males and the age of vaginal opening and first estrus in females. Balano-preputial separation occurred at 29.5 ± 0.3 days in wild-type males, but it was observed at 33.9 ± 0.8 days in *Madin* KO males (Figure 6A). Notably, the weight at balano-preputial separation was comparable between wild-type and *Madin* KO mice (Figure 6B). In females, the age of vaginal opening in wild-type animals was observed at 32.6 ± 0.3 days, but it was delayed at 34.3 ± 0.3 days in *Madin* KO mice (Figure 6C,D). However, the weight at vaginal opening, age, and weight of first estrus was normal in *Madin* KO mice (Figure 6E-H). These results show that sexual maturation is delayed in *Madin* KO mice of both sexes. Accordingly, there was a significant reduction in the delay between the vaginal opening and the first ovulation, indicative of altered timing of puberty onset in female *Madin* KO mice (Figure 6I). During adult life, *Madin* KO mice displayed regular estrous cycles (Figure 6J). Because GnRH neurons are critical regulators of reproductive function (37), we also investigated the distribution of GnRH neurons in the brain of *Madin* KO mice and their wild-type littermates using the iDISCO 3D approach (38). The overall distribution and total number of GnRH neurons were comparable between wild-type and mutant mice (Figure 6K). However, we found fewer GnRH neurons in the olfactory bulb of *Madin* KO mice (Figure 6K).

**Figure 6.**
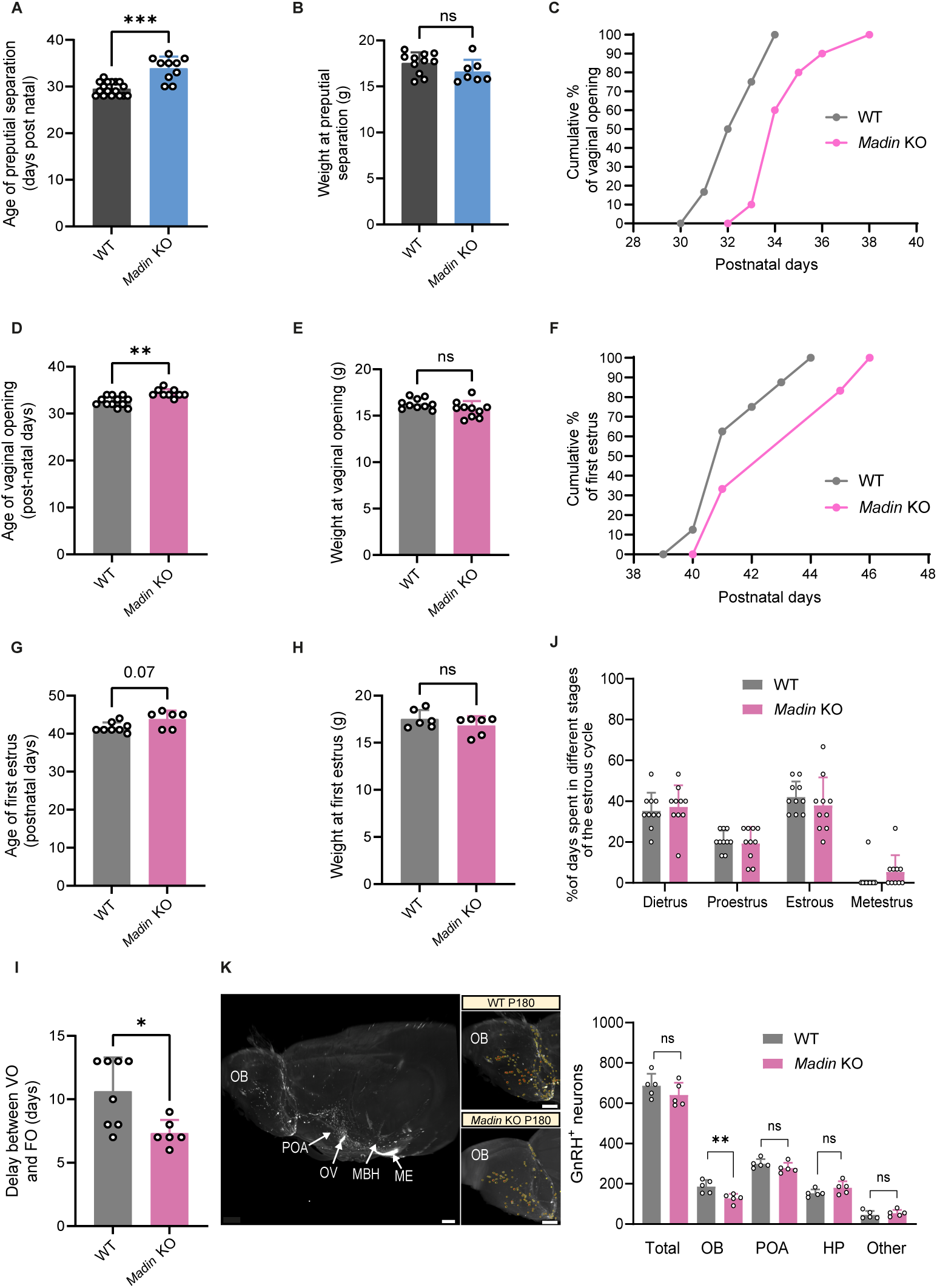
Delayed the onset of puberty in *Madin* KO mice. (**A**) Age and (**B**) weight of balano- preputial separation in male *Madin* KO or wild-type mice (n = 18-20 animals per group). (**C**) Cumulative percentage data for vaginal opening (VO), (**D**) age and (**E**) weight of vaginal opening, (**F**) cumulative percentage data of first estrus, (**G**) age, and (**H**) weight of first estrus, (**I**) delay between VO and first estrus, and (**J**) percentage of days spent in different stages of the estrous cycle in female *Madin* KO or wild-type mice (n = 10-12 animals per group). (**K**) Representative image of cleared brains and immunolabeling for GnRH quantification of the number of GnRH-immunoreactive neurons in female *Madin* KO or wild-type mice at P180 (n = 5 animals per group). Data are presented as mean ± SEM. Statistical significance between groups was determined using a Mann-Whitney Test (A-J). \**\*P ≤ 0.033., **P ≤ 0.002, ***P ≤ 0.0002., ****P ≤ 0.0001.* Scale bar: 500 μm. MBH, mediobasal hypothalamus, ME, median eminence, OB, olfactory bulb, OV, organum vasculosum of the lamila terminalis, POA, preoptic area

### Neuroanatomical alterations in the hypothalamus of *Madin* KO mice

Previous histopathological studies in the hypothalamus of adult patients with PWS reported a significant reduction in the number of OT neurons (39, 40). However, these studies utilized antibodies against the mature form of OT, questioning whether the reduced number of OT- immunopositive neurons is truly due to a loss of OT neurons or caused by perturbations in peptide maturation. Therefore, we used two antibodies targeted against the pro-hormone (*i.e.*, the PS38 antibody) or the intermediate forms of the neurohormone (*i.e.*, the VA10 antibody). We then analyzed the ratio of neurons expressing the intermediate form of OT with those expressing the prohormone and observed an accumulation of the intermediate form of the neuropeptide in *Magel2* KO, *Necdin* KO, and *Madin* KO mice at P0 (Figure 7A).

**Figure 7.**
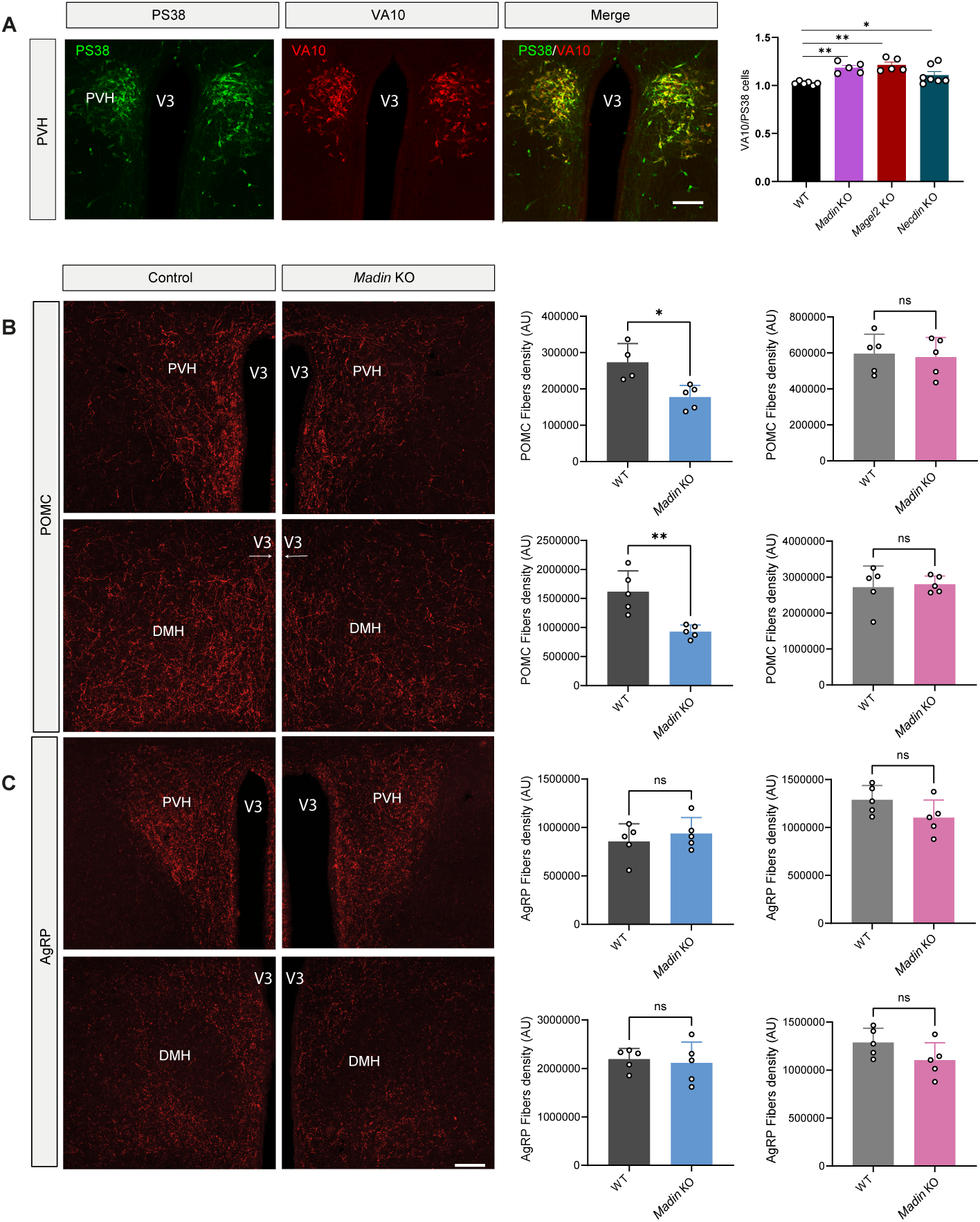
Disruption of hypothalamic melanocortin circuits in *Madin* KO mice. (**A**) Confocal images and quantification of the ratio of PS38- and VA10-immunoreactive neurons in the paraventricular nucleus (PVH) of P0 *Madin* KO and wild-type mice. (**B-C**) Confocal images and quantification of the density of (**B**) POMC- and (**C**) AgRP-immunoreactive fibers innervating the paraventricular and the dorsomedial (DMH) nuclei of the hypothalamus of male (blue) and female (pink) *Madin* KO or wild-type mice at P98 (n = 5 animals per group). Data are presented as mean ± SEM. Statistical significance between groups was determined by a Mann-Whitney test (B-C) or a Mixed-Effect test (A). \**P* ≤ 0.033., \*\**P* ≤ 0.002. Scale bar: 100 μm. V3, third ventricle.

The melanocortin system is a critical component of hypothalamic pathways regulating metabolism. We previously reported a disruption of melanocortin circuits in *Magel2* KO mice (the *Magel2*^tm1Stw^ line) with a reduced density of POMC-immunoreactive fibers in the paraventricular nucleus of *Magel2* KO male mice compared to wild-type mice (19). In contrast, the density of AgRP-orexigenic fibers was normal in *Magel2* KO mice. Consistent with these findings, we found a 1.5-fold and a 1.7-fold reduction in the density of POMC-immunoreactive fibers in the PVH and DMH, respectively, in male but not female *Madin* KO mice compared to wild-type littermates (Figure 7B). The density of AgRP-immunoreactive fibers was normal in the PVH and DMH of male and female mutants (Figure 7C).

### Respiratory alterations in *Madin* KO mice

Respiratory distress was reported in *Necdin* KO neonates and adults (23, 28). The breathing activity of *Madin* KO mice and wild-type littermates was assessed at P30 using *in vivo* whole-body plethysmography (23). During quiet breathing in normocapnia, *Madin* KO mice displayed an increased minute ventilation compared to wild-type mice, mainly due to a tendency for an increased tidal volume, while respiratory frequency was similar in both groups (Figure 8A). There was no difference in the number of apneas and in the irregularity of breathing between wild-type and *Madin* KO mice. A 10 min hypercapnic challenge (4% CO2) increased ventilatory parameters to similar levels in both *Madin* KO and wild-type mice (Figure 8B). There was a tendency for a smaller delta increase in minute ventilation and tidal volume in *Madin* KO mice compared to wild-type mice. During the first 20 min of return to normocapnia following the hypercapnic challenge, the *Madin* KO mice had increased minute ventilation and increased tidal volume compared to wild-type mice, similar to during basal breathing before the hypercapnic challenge (Figure 8C). Also, during this period, *Madin* KO mice showed an overall tendency for more apneas (Figure 8C). This was due to a persistence of the post-hypercapnic increase in apneas throughout the first 20 min of return to normocapnia in mutant mice as wild-type mice showed a reduction in their number of apneas starting ∼5 min after the return to normocapnia (Figure 8D). These results indicate an increased ventilation of *Madin* KO mice in basal normocapnic condition, and a persistent increase in apneas following a hypercapnic challenge.

**Figure 8.**
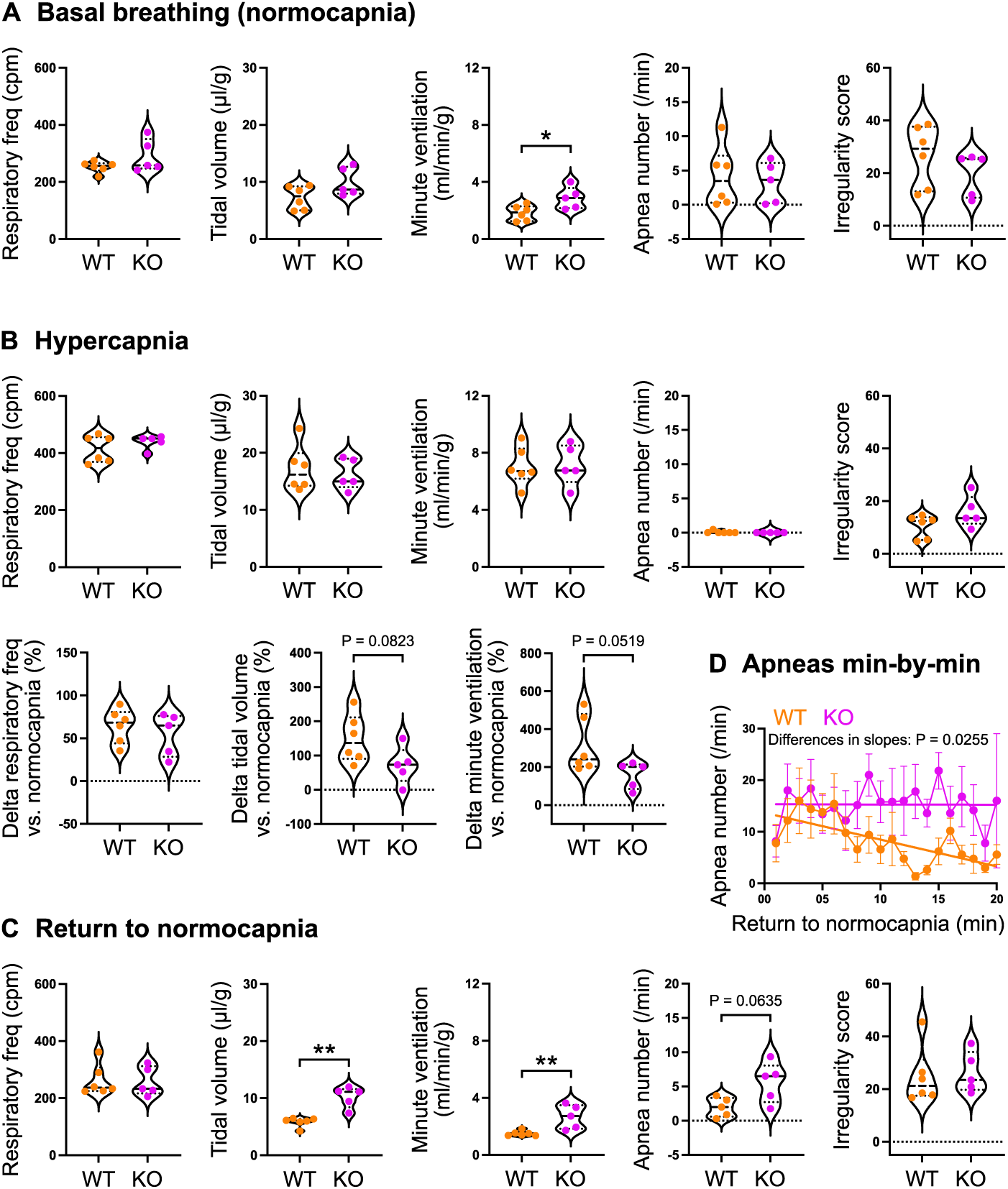
Increased ventilation and post-hypercapnic apneas in *Madin* KO mice. (**A-C**) Analysis of the breathing activity in P30 *Madin* KO (n = 5 animals per group) and WT (n = 6 animals per group) mice using *in vivo* whole-body plethysmography (**A**) during basal conditions (normocapnia) and (**B**) during a 10 min hypercapnic challenge (4% CO_2_) and (**C**) during the first 20 min of return to normocapnia following the hypercapnic challenge. (**D**) Analysis of the number of apneas per minute during the first 20 min of return to normocapnia. Statistical significance between groups was determined by a Mann-Whitney test (A-C) or a Linear fitting with the least squares regression method, extra sum-of-squares F Test to compare slope differences (D). \**P* < 0.05., \*\**P* < 0.001. cpm, cycles per minute.

### Hypothalamic genes are differentially expressed in *Madin* KO mice during the circadian cycle

Transcriptional studies have proven helpful in understanding disease mechanisms. Therefore, we performed bulk RNA sequencing on the hypothalamus of adult *Madin* KO mice and their wild-type littermates. Since *Magel2* and *Necdin* have a circadian expression, with a significant ∼1.5-fold increase in expression level during the night versus day (Figure 9A), and it has been shown dysregulation of the circadian activity and circadian genes (26, 41) in single *Necdin* and *Magel2* KO, we investigated genes that were differentially expressed between the middle of day (ZT6) and the middle of the night (ZT18) in the hypothalamus of *Madin* KO versus wild-type littermates. In the hypothalamus of wild-type mice, we found 683 genes that were differentially expressed between the ZT6 and ZT18 (Figure 9B). It may be assumed that these genes display a rhythmic pattern over the nychthemeral cycle. Remarkably, 681 out of these 683 genes lose their differential expression between day and night in the *Madin* KO mice (Figure 9B,C). Because measurements were taken at only two time points within the nychthemeral cycle, it was not possible to determine whether these genes became arrhythmic or if their phase had been shifted. Nevertheless, a Panther analysis revealed that in this list of 681 genes that displayed a differential day/night expression only in wild-type mice, the terms ‘‘nervous system development”, “cell differentiation”, and “cell-cell adhesion” were among the most significant enriched annotations in Gene Ontology biological processes (Figure 9C). We also identified 78 genes in the hypothalamus of *Madin* KO mice that acquired a day-night difference that was not found in wild-type mice (Figure 9A-B). As previously highlighted, it cannot be determined whether these genes became rhythmic when they were not in the wild-type mice, or if a phase shift enables the detection of a day/night difference that was undetectable with two measurement times in the wild-type. Overall, it appeared that 99% of genes with a differential day/night expression in both wild-type and *Madin* KO mice were dependent on *Magel2/Necdin* expression. These genes could be classified into three different categories depending on whether 1) they lost their differential expression, 2) they had a modified rhythmic pattern, or 3) they acquired a differential day/night expression (Figure 9 B-C).

**Figure 9.**
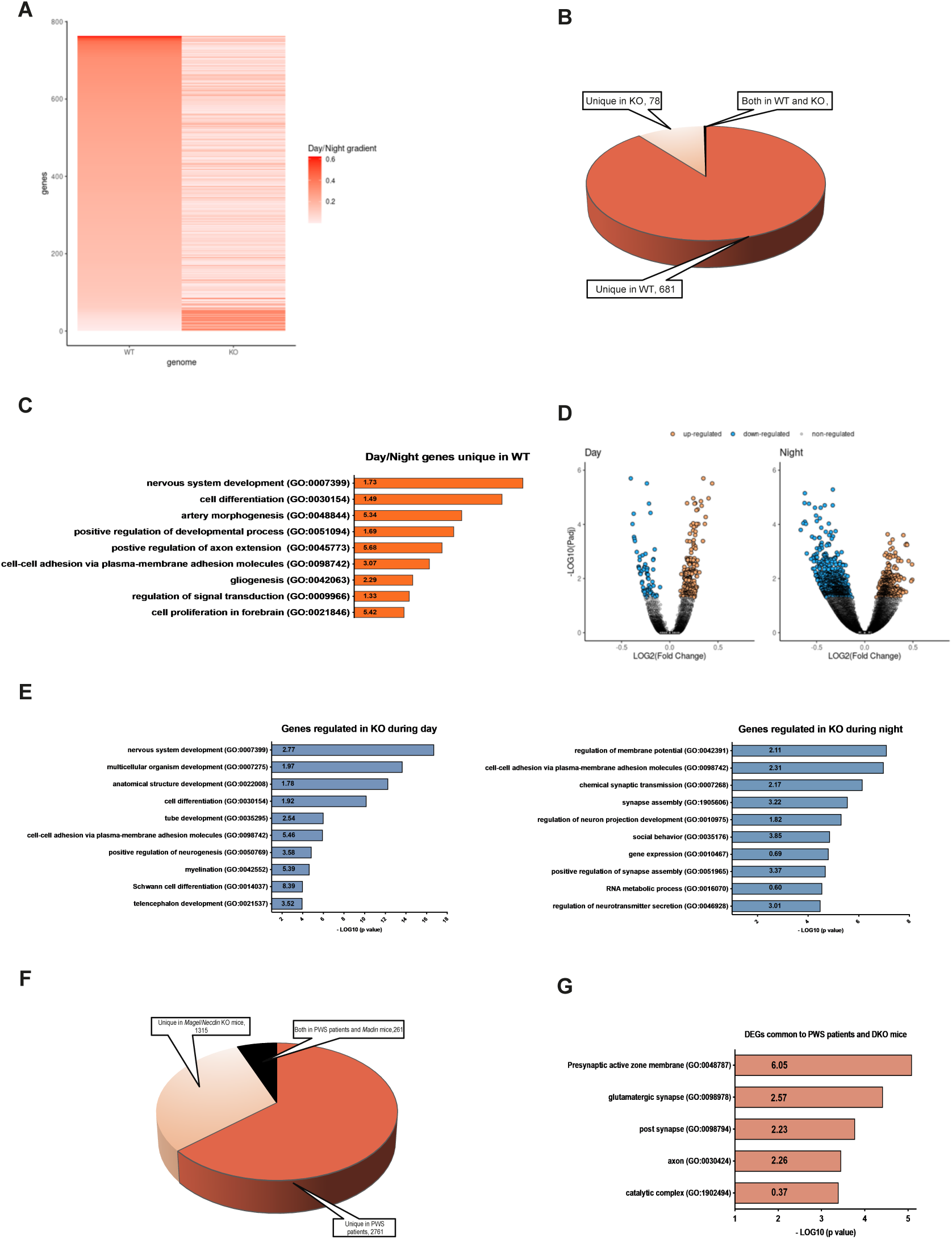
Transcriptomic analysis of genes that are dysregulated in the hypothalamus of *Madin* KO mice. (**A**) Heat maps representing the gradient of change (as the absolute value of Log2 fold change) between day and night for all expressed genes in wild-type (WT, left part) and for corresponding genes displayed in the same order in *Madin* KO mice (right part) at P70- P98 (n = 5 animals per group). (**B**) Impact of *Magel2*+*Necdin* double deletion on differentially day/night expressed genes. 681 genes differentially expressed between day and night in WT lose their differential day/night expression in KO (Unique in WT, 90%), while 78 genes acquire a *de novo* day/night difference in expression (Unique in KO, 10%). Only two genes display a differential day/night expression in both mouse lineages (Both in WT and KO). (**C**) Functional characterization by Panther Analysis of genes belonging to the category “Unique in WT” showing the top 10 Gene Ontology biological processes with the highest *p* value. The numbers inside the columns correspond to the fold enrichment. (**D**) Volcano plots illustrating up-regulated (orange points) and down-regulated (blue points) genes in *Madin* KO *versus* WT mice during the day (left part) and during the night (right part). During the day, 74% of genes are up-regulated in *Madin* KO mice while 68% are down-regulated during night. (**E**) Functional characterization by Panther Analysis of genes impacted in *Madin* KO mice during the day (left part) and during the night (right part) showing the top 10 Gene Ontology biological processes with the highest *p* value. The numbers inside the columns correspond to the fold enrichment. (**F**) Diagram showing genes that are commonly dysregulated in the hypothalamus of *Madin* KO mice and patients with PWS (39), genes that are uniquely dysregulated in *Madin* KO mice, and genes that are uniquely dysregulated in patients with PWS. (**G**) Functional characterization by Panther Analysis of genes commonly impacted in the hypothalamus of *Madin* KO mice and PWS patients showing the top 5 Gene Ontology biological processes with the highest *p* value.

We further analyzed the impact of the combined deletion of *Magel2* and *Necdin* on hypothalamic gene expression during the day (ZT6) and the night (ZT18). During the day, 261 genes had a different expression level in *Madin* KO compared to wild-type mice, 68 (26%) being down-regulated and 193 (74%) up-regulated (Figure 9D). As shown after Panther analysis, these 261 genes impacted by *Magel2*/*Necdin* deletion were shown to be significantly associated with different functions such as “development”, “differentiation”, and “myelination” (Figure 9E, left part). At night, more genes (1295 genes) had their expression level differentially regulated in *Madin* KO mice, but while 874 (68%) of these genes displayed a down-regulation, 421 genes (32%) were up-regulated (Figure 9E, right part). These 1295 regulated genes were mostly associated with synaptic transmission, cell-cell adhesion and social behavior (Figure 9E). Among all regulated genes either during the day or during the night (1556 genes), 258 were found to display a day/night differential expression in wild-type meaning that around 17% of genes whose expression was altered in *Madin* KO mice could be circadian rhythmic genes (data not shown).

We also explored whether there were genes that were commonly dysregulated in *Madin* KO mice and patients with PWS by comparing our transcriptomic data with an RNA sequencing analysis previously performed in the hypothalamus of patients with PWS (39). We found that 261 genes were commonly dysregulated in both *Madin* KO mice and PWS patients (Figure 9F and Supplementary Table 1). 1315 genes were dysregulated only in *Madin* KO mice, and 2761 were dysregulated only in patients with PWS (Figure 9F). A Panther analysis revealed that in this list of 261 genes that were commonly dysregulated in mice and humans, the terms ‘‘presynaptic active zone membrane”, “glutamatergic synapse”, “post synapse”, and “axon” were among the most significant enriched annotations in Gene Ontology biological processes (Figure 9G).

## Discussion

The present study describes the phenotypic characterization of a novel mouse model with a deletion of both *Magel2* and *Necdin,* two genes also being deleted in PWS. The rationale for generating this mouse model was to avoid a potential functional redundancy between these two genes that could mask some symptoms in the *Necdin*- or *Magel2*- single KO mice, both genes encoding MAGE proteins that bind E-3 ubiquitin ligases specifying proteins for ubiquitination (42). Firstly, we confirmed that *Magel2* and *Necdin* are co-expressed in many brain nuclei throughout life and co-regulated, potentially sharing a common enhancer. More importantly, we showed that when *Magel2* is deleted, *Necdin* expression is increased and *vice versa*. Such overexpression could induce a compensatory mechanism, but *Necdin* or *Magel2* overexpression could also be responsible for part of the phenotype in *Magel2* KO or *Necdin* KO mice, respectively. Indeed, it has been reported that a twofold increase in *Necdin* induced ASD-related behaviors (43). Similarly, overexpression of the N-terminal region of Magel2 is lethal at embryonic or neonatal stages, whereas normal production of this Magel2 protein is not lethal, indicating the toxic effects of the overexpression of the protein (44). Thus, the *Madin* KO mouse model with a deletion including both genes is genetically and phenotypically more reliable and relevant to study PWS since both genes are deleted and the compensatory mechanisms and/or phenotypes linked to overexpression that could occur in single KO (*i.e*., in *Magel2* or *Necdin* KO mice) do not occur in *Madin* KO mice, similar to the situation in PWS.

Our teams have created and extensively studied the *Necdin ^tm1.1Mus^*and *Magel2^tm1.1Mus^* KO mouse models from which *Madin* mice have been created. We and others, such as the R. Wevrick’s team, have revealed important and specific roles for each of these genes in the pathophysiology of PWS, and these mouse models have been used in several pre-clinical studies. However, phenotypic differences were observed between the six different *Necdin* KO models and the three different *Magel2* KO models that have been described (Table 1 and (8)). Those differences might be explained by the genomic constructions causing either a deletion and lack of transcripts or the creation of a truncated or fused protein that might induce an additional phenotype to the loss of function of the normal proteins. Furthermore, at least in *Necdin ^tm1.1Mus^*and *Magel2^tm1.1Mus^* KO mice, a stochastic and very low expression of the maternal allele has been detected for *Necdin* (34) and *Magel2* (28), respectively. This weak expression was sufficient to alleviate the phenotypes (34) and explained part of the variability between pups from the same litter. Similar weak expression was detected in patients with PWS and might also explain part of the variability in the phenotype (34).

**Table 1.**
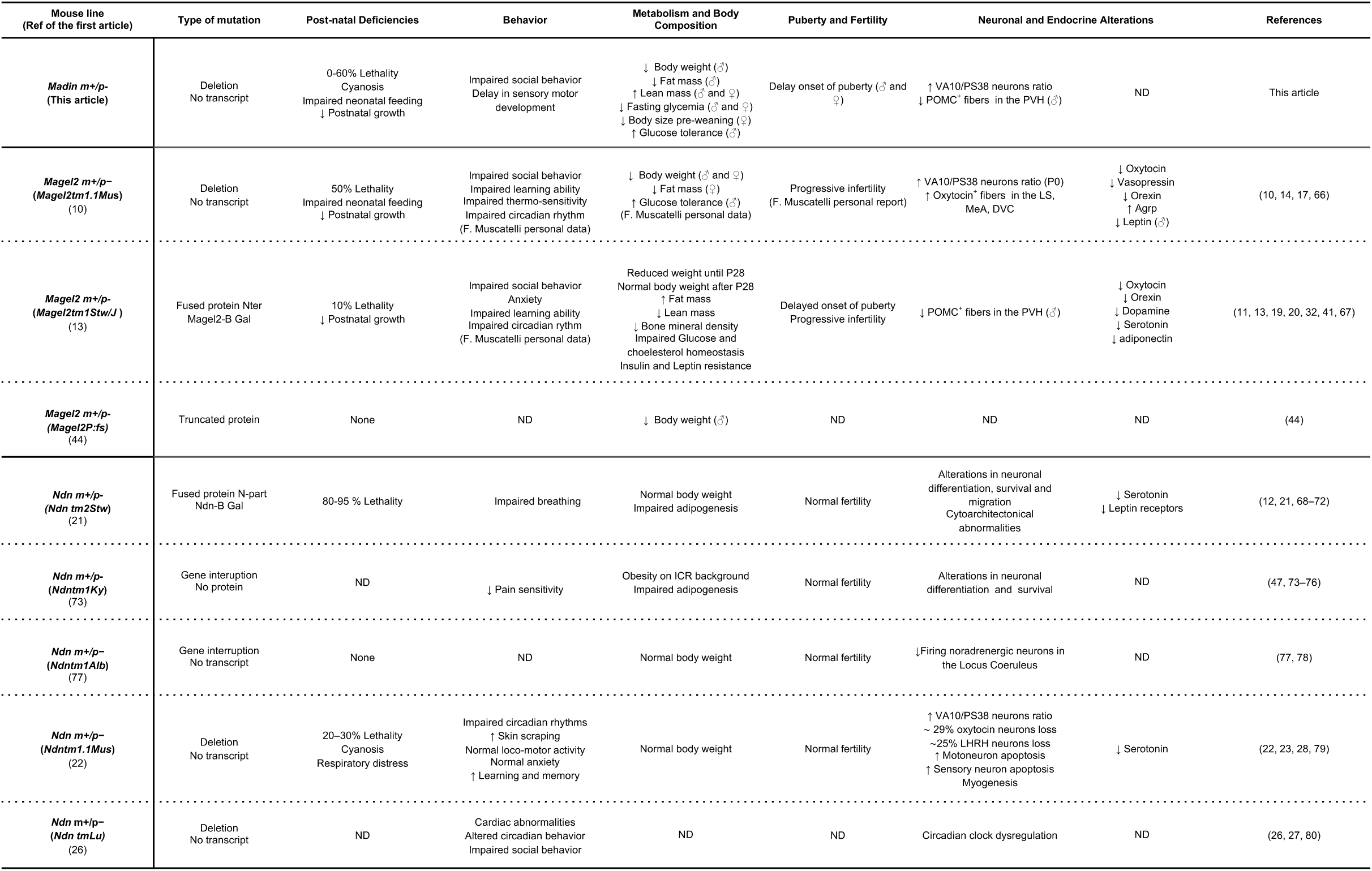
Phenotypic comparison of *Madin* KO mice with *Magel2* and *Necdin* single KO models. Up- and down-regulation of physiological parameters are represented by arrows (↑ and ↓). Data from *Magel2* and *Necdin* KO models are derived from (8).

In the present study, we produced several cohorts of *Madin* KO mice in different animal facilities, all on the same C57Bl6/J genetic background, and observed variability in the severity of the phenotype (in particular on the mortality rate at birth) linked to the cohort. We also observed this variability with the *Magel2^tm1.1Mus^*mice (unpublished data). We could correlate the phenotypic variability of *Madin* KO mice with the different levels of sanitary status of the animal facilities in which animals were housed, *i.e.*, in terms of pathogens and sterility of food and water. For example, when animals were housed in a pathogen-free animal facility and fed with sterile food, the phenotype was much less severe than when they were housed in a conventional animal facility. It is interesting to note that MAGE genes, conserved in all eukaryotes, play a role in adaptation to stress (*e.g.*, nutritional, genotoxic, and heat) (42) and that in the absence of stress, their function is not revealed. It is therefore likely that the phenotype of *Madin* KO mice could be highly dependent on the environment (including pathogens, food and water) and this issue should be taken into consideration in all animal houses, where the absence of pathogens does not reflect real life. The environment is also a challenge in patients with PWS. For example, respiratory failure is the most common cause of death (73% of infants) and always occurs in a stressful environment (*e.g*., airway infection, hypercapnia, hypoxia). This observation reveals the infants’ inability to adapt their respiratory response (45).

Feeding difficulties, respiratory deficits, and low body weight are among the first symptoms observed in babies with PWS. These early phenotypic traits are also found in our *Madin* KO neonates that lacked milk in their stomachs, appeared cyanotic, and displayed a low body weight and somatic growth from birth to weaning. The absence of milk in the stomach was also observed in *Magel2 ^tm1.1Mus^* KO pups and was shown to result from a suckling deficit caused by a defect in OT maturation and release. A similar reduction in the number of OT neurons was reported in the *Ndn^tm1-Mus^* KO mice (22). Interestingly, we also found a defect in OT maturation in *Madin* KO pups. The cyanotic appearance was also observed in *Necdin ^tm1.1Mus^* KO neonates. It reflects respiratory distress that was extensively studied in *Necdin ^tm1.1Mus^* KO mice and is characterized by frequent apneas, irregular rhythms in basal conditions, and altered ventilatory response to hypercapnia (23). We did not observed cyanosis in *Madin KO* pups born in a free- pathogen animal facility, but during adult life we detected differences in ventilatory parameters in normal conditions and also an abnormal persistence of apneas after an hypercapnic challenge, revealing alleviated respiratory deficits.

One of the hallmarks of PWS is the progression of symptoms over time. The motor, and possibly sensory deficits underlying hypotonicity of infants with PWS tend to disappear later in childhood. We did not observe motor deficits in *Madin* KO pups but our results suggest a delay in maturation of touch/proprioception sensory modalities. PWS is also characterized by endocrinopathies including growth deficiency, hypogonadism and delayed puberty onset which we also observed in our *Madin KO* mouse model. The feeding difficulties found in infants with PWS are replaced by hyperphagia, leading to the development of obesity (46). Despite the development of many rodent models of PWS, only two models have somewhat reproduced a phenotypic trait related to weight gain. *Magel2* KO mice do not become obese but tend to gain more weight after weaning (11), and a strain of *Necdin* KO mice (on ICR background) (47) become obese on a high-fat diet. More conflicting findings have been reported with the feeding behavior of *Magel2* KO mice with some studies describing moderate increases (48) and others slight decreases (11, 13) in food intake in mutant mice. All the other mouse models of PWS tend to be leaner than control littermates (8). Similarly, male *Madin* KO mice have reduced body weight and fat mass, and normal food consumption, energy expenditure, and locomotor activity. In addition, male *Madin* KO mice display an improved glucose tolerance similar to the ameliorated glucose metabolism generally observed in patients with PWS (36). Altogether, these findings suggest that several metabolic and feeding behavior defects observed in PWS might partly be independent of *Magel2* and *Necdin*.

Behavioural changes are present throughout the patient’s life. Anxiety, interactions with novel objects or social interactions are altered in *Necdin* (27) and *Magel2* KO mice (14, 17), with a variability in the severity of the phenotype, partly due to the environmental context. *Madin* KO mice showed a preference for the novel object *versus* a familiar one and for a congener mouse *versus* an empty cage. However, the interest in moving towards the novel object or the time spent to socialize is reduced compared to the controls. Importantly, the cohort of *Madin* KO mice investigated for adult behavior was generated in a “sterile” context (associated with 0% lethality), which could explain why we observed a milder alteration of social behavior compared with *Magel2* mice. Intriguingly, recent studies have implicated alterations in GnRH number and function, including during early postnatal development, in adult social interaction and cognition (38, 49, 50). *Madin* KO mice show a selective reduction in the number of GnRH cell bodies in the olfactory bulb, a neuronal population recently shown to be involved in processing socially-relevant odors required for congener recognition (51). This finding provides a potential link between GnRH neuron alterations and social behavior deficits observed in *Madin* KO mice. The control and maintenance of GnRH production in the brain after birth could this be an interesting actionable target for managing cognition and social interactions in PWS. Similar approaches have shown promise in other chromosomal disorders such as trisomy 21 (50).

Because *Magel2* and *Necdin* have a circadian expression (26), transcriptomic analyses were performed on hypothalami collected in the middle of the day (ZT6) and in the middle of the night (ZT18) to identify alterations in rhythmic patterns of gene expression. We found that the day/night differential expression of the master core-clock gene *Arntl* (also known as *Bmal1*) was blunted in *Madin* KO mice. While a phase shift in its circadian pattern cannot be excluded, its decreased expression level at night is consistent with the reduction in BMAL1 protein expression observed after *Necdin* repression in U2OS cells (26). It has been previously shown that *Necdin* regulates BMAL1 stability and the amplitude of the rhythm of several core-clock genes such as *Bhlhe40* (Dec1) and *Bhlhe41* (Dec2) (26). Accordingly, some core-clock genes other than *Bmal1*, such as *Npas2*, *Nr1d1* (Rev-erbα), *Nr1d2* (Rev-erbβ), *Bhlhe40*, and *Bhlhe41*, were also affected in *Madin* KO mice. These genes either lost their Day/Night differential expression (*Bhlhe40, Nr1d1,* and Nr1d2), exhibited altered levels at night (*Npas2* and *Bhlhe41*), or, similar to *Arntl*, displayed both alterations (*Nr1d1*). We found that genes affected in *Madin* mice were significantly overrepresented in the category “rhythmic process” with twenty-one identified genes (data not shown). In addition to the core-clock genes mentioned above, other genes in the 21-gene list are histone-modifying enzymes such as *Crebbp, Ep300,* or *Kmt2a*, known to play a crucial role in circadian-clock-output gene expression by contributing to the rhythmic recruitment of the CLOCK-BMAL1 transcription factor complex to circadian gene promoters (52). Some genes regulated in the *Madin* KO mice could also serve as hubs between core-clock genes and circadian-clock-output genes.

Previous transcriptomic analyses performed on *post-mortem* hypothalamic tissues of patients with PWS identified changes in molecular pathways involved in neuronal loss, neuroplasticity, and neuroinflammation (39). Interestingly, we found 261 genes that were commonly dysregulated in both *Madin* KO mice and patients with PWS, and the functions of these genes appeared to be related to synaptic and axonal function, making the *Madin* KO mice a particularly relevant model to study neurodevelopmental and neurocircuits defects associated with PWS. In support of the alteration of genes involved in axonogenesis, we observed structural alterations in POMC neuronal circuits in *Madin* KO mice, similar to what we previously reported in the *Magel2* KO model (19).

In conclusion, we observed a wide range of phenotypic traits in *Madin* KO mice that cover neurodevelopmental and behavioral symptoms of PWS, recapitulating part of the phenotypes observed in *Necdin* and *Magel2* KO mice. It would be important to compare the phenotype of *Magel2* and *Necdin* simple KO with that of *Madin* KO mice in the same experimental and environmental conditions to rigorously determine whether the phenotype of *Madin* KO mice is the addition of the *Magel2* and *Necdin* simple KO phenotypes only or whether it results from a more complex phenotype. Nevertheless, from a genetic point of view and because of the interplay between *Magel2* and *Necdin*, *Madin* KO mice are currently the most appropriate and relevant pre-clinical model for studying PWS.

## Methods

### Sex as a biological variable

Our study examined male and female animals. Both sexes were pooled when no sex differences were found.

### Study approval

All experiments were performed in accordance with the Guide for the Care and Use of Laboratory Animals (N.R.C., 1996) and the European Communities Council Directive of September 22th 2010 (2010/63/EU, 74) and the approved protocol (APAFIS# 13387– 2017122712209790 for the studies performed in Lille and accreditation no. B13-055-19 for the studies conducted in Marseille) by the Ethical Committee of the French Ministry of Agriculture.

### Trans-allelic recombination to generate the *Madin* KO mouse model and housing conditions

We used a Cre-loxP site-specific recombination to mediate efficient trans-allelic recombination *in vivo* (53) between loxP site in the *Necdin* KO mouse (*Ndn^tm1-Mus^*) (22) and loxP site in the *Magel2* KO mouse (*Magel2^tm1-Mus^*) (10). This approach can be used because both loxP sites are oriented in the same direction allowing the creation of one allele with the deletion of the DNA region between both loxP sites. We used an *Hprt-*Cre driver mouse in which a strong CAG promoter is inserted into the X-linked *Hprt* locus allowing a strong, constitutively expressed, Cre recombinase. *Hprt*-Cre/*Magel2* KO (-m/+p) male mice were generated and then crossed with Heterozygous *Necdin* KO female mice (-m/+p).

### Behavioral tests in pups

Behavior tests performed during the two first weeks of postnatal life followed a protocol described in (35). The age of eye and auditory canal opening was checked by visual inspection. The righting response was performed by scoring the time for righting, *i.e.*, the ability of the pup put on its back to stand up on its paws (scoring the time for righting). The rooting response assessed the disappearance of the rooting reflex triggered by bilateral stimulation of the snout. 3 consecutive tests were performed and scored, with the total score plotted on a graph (Score 0: pup roots; score 1: pup stops to root; score 2: pup removes the head from the fingers). The paw position test is indicative of the appearance of an adult pattern when all four paws were placed flat on the ground (score 1 per paw placed flat on the ground). The bar holding test consisted in scoring the time during which the pup is able to hang when putting its ventral face on the bar just at the level of the diaphragm, the position was unstable and the pup hanged onto the bar to prevent falling. The pulling up on the bar after hanging test consisted in performing the bar holding test and assessing the pulling up on the bar after hanging. Three scores were given: score 1: the pup pulled up by drawing the bar towards him; score 2: the pup braced himself with his hind feet; score 3: the pup stood or walked on the bar. The climb the slope test consisted of the pup climbing up a 30° slope. The pup is positioned with the head downward (stay still), it turned the head, then the body at 90°, then at 180° and climbed the slope. The response is due to the maturation of semicircular canals (inner ears). In the vertical climbing test, the pup climbed along the wire mesh of vertical grid. The score was the number of mesh wires that the pup covers in 20 sec. In the cliff drop avoidance test, the pup should turned away from the edge of a step, starting with the forepaws dangling over the edge of the slice (score 0: it remained motionless or fell from the edge; score 1: it turned the head aside; score 2: it turned the head aside, pulled back the forepaws from the cliff and put them on the slice; score 3: it initiated a backward movement.

### Behavioral studies in adult mice

Behavioral tests in adulthood were performed by Phenotype Expertise, Inc. (France) with an expert behaviorist. The number of tested animals was based on previous publications and Phenotype Expertise experience. For all tests, mice were first acclimated to the behavioral room for 30 minutes.

#### Rotarod

Mice were challenged to walk on a rotating rod that increases in speed over a predetermined period of time (for 5 min) until the animal falls. The latency to fall from the rod was determined and taken as a measure of motor function. The test was performed over three successive days. The daily trial consisted of three measures spaced by 5 minutes, a mean of the three measures was considered for each day.

#### Elevated plus maze

The device consists of a labyrinth of 4 arms of 5 cm width located 80 cm above the ground. Two opposite arms are open (without wall) while the other two arms are closed by side walls. The light intensity was adjusted to 20 Lux on the open arms. Mice were initially placed on the central platform and left free to explore the cross-shaped labyrinth for 5 minutes. The maze was cleaned and wiped with H_2_O and with 70% ethanol between each mouse. Animal movement was video-tracked using Ethovision software 11.5 (Noldus). Time spent in open and closed arms, the number of entries in open arms, as well as the distance covered, are directly measured by the software.

#### Open field

The open field test was performed in a 40 x 40 cm square arena with an indirect illumination of 60 lux. Mouse movement was video-tracked using Ethovision software 11.5 (Noldus) for 10 minutes. Total distance traveled and time in center (exclusion of a 5 cm border arena) are directly measured by the software. Grooming (time and events) and rearing were manually counted in live using manual functions of the software, by an experimented behaviorist. The open-field arena was cleaned and wiped with H_2_O and with 70% ethanol between each mouse.

#### Spontaneous Social Interaction in an open field arena

Animals were first habituated to the apparatus for 30 min. Social interaction was measured using pairs of mice from different housing cages, and having the same genotype, the same gender, and approximately the same body weight. Each pair was placed in the open field arena for 15 min during which different behavioral parameters reflecting social interaction between the two mice were recorded (following and sniffing). A social interaction index was defined as the percentage of counts and duration of social events over the total social and individual events.

#### New object recognition

The arena used for the novel object recognition test was the same used for the open-field test. The arena was cleaned and wiped with 70% ethanol between each mouse. Two identical objects (50 ml orange corning tube) were placed in the opposite corners of the arena, 10 cm from the side walls. The tested mouse was placed at the opposite side of the arena and allowed to explore the arena for 10 min. After 1h, one object was randomly replaced with another novel object, which was of similar size but differ in the shape and color with the previous object (white and blue lego bricks), the other object (same object) was kept. Then, the same mouse was placed in the arena and allowed to explore the two objects (a new and an "old" familiar object) for 10 min. The movement of the mice was video-tracked with Ethovision 11.5 software. Time of exploration of both objects (nose located in a 2 cm area around object) was automatically measured by the software. The traveled distance to reach the novel object or the same object (old object) was measured.

#### Three-chamber social preference test

The three-chamber apparatus consisted of a Plexiglas box (50x25 cm) with removable floor and partitions dividing the box into three chambers with 5-cm openings between chambers. The task was carried out in four trials. The three-chambers apparatus was cleaned and wiped with 70% ethanol between each trial and each three-chamber test experiments. In the first trial (habituation), a test mouse was placed in the center of the three-chamber unit, where two empty wire cages were placed in the left and right chambers to habituate the test mouse to arena. The mouse was allowed to freely explore each chamber. The mouse was video-tracked for 5 min using Ethovision software. At the end of the trial, the animal was gently directed to the central chamber with doors closed. In the second trial (social exploration), an 8 weeks old C57BL/6J congener mouse (S1) was placed randomly in one of the two wire cages to avoid a place preference. The second wire cage remained empty (E). Then, doors between chambers were opened and the test mouse was allowed to freely explore the arena for 10 min. The measure of the real social contact is represented by the time spent in nose-to-nose interactions with the mouse. This test was performed using grouped-house mice.

### Metabolic phenotyping

Animals were weighed daily from birth to weaning (P21) and weekly from P21 to P168 using an analytical balance. Body length was measured at P24 and P60 with a rigid metric ruler. Body composition analysis was conducted weekly from P25 to P168 using a Minispec LF Series (Bruker Corporation, Massachusetts). Fat mass, lean mass, and free body fluids measurements were expressed as a percentage of total body weight. Infrared pictures were taken with a hand-held camera (E8, with an accuracy of 2% max 2 °C, FLIR) on freely moving and unshaven mice at P120 to assess interscapular BAT temperature. Food intake, O_2_ consumption and CO_2_ production, energy expenditure, respiratory exchange ratio (*i.e.*, VCO_2_/O2), and locomotor activity (X and Y axis) were monitored in fed mice, fasted mice for 24-hours and after a 48-hour refeeding period at P180 using a combined indirect calorimetry system (TSE Pheno Master Systems GmBH, Germany). The mice were acclimated in monitoring chambers for two days, and the data were collected for 6 days. These physiological measures were performed at the Mouse metabolic phenotypic platform of the University of Lille. Glucose tolerance tests were conducted in mice at ∼P130 of age through i.p. injection of glucose (2 g/g body weight) after 6 hours of fasting. Blood glucose levels were measured at 0, 15, 30, 60, 90, 120, and 150 minutes post-injection, as previously described (54).

### Reproductive Phenotyping

Starting on P21, female mice were inspected daily for imperforation of the vaginal membrane (“vaginal opening”, VO). After that, vaginal smears were collected daily and analyzed under the microscope to identify the onset of puberty (first appearance of two consecutive days where vaginal smears contained cornified cells) and the specific day of the estrous cycle, as described previously (55, 56). Male mice were checked daily for balano-preputial separation as an external sign of puberty onset.

### Hormone assays

Serum insulin levels were measured in P130 mice before, 15, and 30 minutes after intraperitoneal i.p. glucose administration (2 g glucose/kg body weight) using commercially available insulin ELISA kits (Mercodia). Serum leptin levels were measured in fed mice at P160-P240 using commercially available leptin ELISA kits (DuoSet Ancillary).

### *In vivo* plethysmography recordings

Unrestrained, non-anesthetized mice were monitored at P30 for their breathing activity using whole-body plethysmography equipment (EMKA Technologies, Paris, France). A constant air flow (0.5 l/min) circulated inside the 200 ml plethysmography chambers (maintained at 25 ± 0.5 °C) using a vacuum pump (Vent 4, EMKA, Paris, France). Calibration of the amplitude of the signal was ensured by injecting 1 ml of air into the chamber during each recording session. Analog signals were acquired through a usbAMP device and processed using EMKA technologies IOX software (EMKA Technologies, Paris, France). Respiratory parameters (frequency, tidal volume, minute ventilation, apneas and irregularity score) were analysed using the Spike2 software (Cambridge Electrical Design, Cambridge, UK). Apneas were defined as a prolonged expiratory time equivalent to the loss of two respiratory cycles. The day before the experiment, wild-type and *Madin* KO littermates were habituated in the plethysmography chambers for 2 hours to reduce stress effect on breathing. Respiratory responses to hypercapnia were then evaluated by recording breathing activity for 1 hour under normocapnia, followed by a 10-minute hypercapnic challenge (4% CO2), and a subsequent 30-minute recovery period under normocapnia. Measurements were performed during quiet breathing pre- and post-challenge, and during the last 5 minutes of the challenge. Apneas were also quantified on a minute-by-minute basis during the post-challenge period.

### Neuroanatomical studies

For standard immunohistochemistry, paraformaldehyde-fixed brains were then frozen, sectioned at 30-µm thickness, and processed for immunofluorescence using standard procedures (57). Brain sections were mounted on glycerol-based medium with DAPI to visualize cell nuclei. For the iDISCO+ immunolabeling, whole brains were first dehydrated, then incubated with the primary and secondary antibodies and cleared as previously described (58). The primary antibodies used for IHC were as follows: rabbit anti-POMC (1:10,000, Ref# H-029-30, Phoenix Pharmaceuticals), rabbit anti-AgRP (1:1,000, Ref# H-003-53, Phoenix Pharmaceuticals), mouse anti-neurophysin 2 clone PS38 (1:1000, Ref# MABN844, Merck Millipore), rabbit anti-GnRH (1:3,000, Ref# #26950-1-AP, Proteintech), and rabbit anti-Necdin (1:500, Ref# 07-565, Merck Millipore). The primary antibodies were visualized with goat anti-rabbit IgG conjugated with Alexa Fluor 568 (1:500, Ref# A11011, Invitrogen) or donkey anti-rabbit IgG with Alexa Fluor 568 (1:1,000, Ref# A10042, Invitrogen) or anti-mouse IgG conjugated with Alexa Fluor 647 secondary antibodies (1:500, Ref# A21235, Life Technologies).

2D images were acquired using a Leica Stellaris 5 confocal Microscope equipped with a 20x objective. Quantifications were performed in two sections per animal. Slides were numerically coded to obscure the experimental group. The image analysis was performed using the Fiji software (NIH) as previously described (57). For the quantitative analysis of fiber density (for POMC and AgRP), a maximum intensity projection was performed on 5 µm of the Z-stack. The threshold was set manually to ensure that only a positive signal was measured. Images were then binarized, and a standardized region of interest (ROI) was placed within the nucleus of interest. The software then calculated the number of pixels in the ROI corresponding to the signal of interest. This pixel count was finally normalized to the dimensions of the ROI, ensuring comparability across all images. The integrated intensity, which reflects the total number of pixels in the binarized image, was then calculated within the ROI of each image of the stack (58, 59) (58, 59). For the quantitative analysis of cell numbers, PS38-immunopositive cells were manually counted using the Fiji software. Only cells with corresponding DAPI-stained nuclei were included in our counts.

3D imaging was performed on a light sheet Ultramicroscope I (LaVision BioTec) equipped with a 1.1X/0.1NA objective and an Andor Neo 5.5 sCMOS camera. ImspectorPro software (LaVision BioTec) was used for image acquisition, and the z-step between each image was fixed at 5 μm. Image stacks were converted to Imaris files (.ims) using ImarisFileConverter, and 3D reconstruction was performed using “volume rendering” of Imaris 9.8 (Oxford Instruments). Automatic spot detector function of Imaris software was used to count the number of GnRH immunoreactive cell bodies. Anatomic 3D neurons’ distribution was done manually.

### *In situ* hybridization

Brain sections from embyros were post-fixed with 4% paraformaldehyde and processed for *in situ* hybridization using antisense digoxigenin-labeled *Necdin* and *Magel2* riboprobes as previously described (9). *In situ* hybridization on adult brain sections were performed using fluorescent labelling as previously described (34). The *Ndn* riboprobe (290 bp) hybridized to the 3’ UTR of the *Ndn* mRNA (nt 2130 to 2420; accession number D76440). The *Magel2* riboprobe (318 bp) hybridized to the 3’ part of the Magel2 ORF (nt 3730 to 4048; accession number AK086725).

### RNA extraction and RT-qPCR analyses

Wild-type and mutant newborns were sacrificed at P0. The hypothalamus was quickly dissected on ice and rapidly frozen in liquid nitrogen, then stored at −80 °C. Total RNA was isolated using the RNeasy® Mini Kit (Qiagen, cat #74104), according to the manufacturer’s protocol and cDNAs were obtained by reverse transcription using QuantiTect® Reverse Transcription Kit (Qiagen, cat #205311), starting with 600 ng of total RNA. We performed RT-qPCR using Sybrgreen-based application (on the LightCycler^®^480 Instrument) to measure mRNAs quantity of *Necdin* and *Magel2* transcipts normalized with *GapdH* and *Actin* reference transcripts as previously described (61). The qPCR primers used were the following: *Necdin*, Forward :AACAACCGTATGCCCATGA and Reverse : CTTCACATAGATGAGGCTCAGGAT; *Magel2*, Forward : CTGGGAGATTCAGAGGGCTA/ and Reverse : TGCGGAGTGTAGAGGGATTC.

### RNA expression analysis by RNA sequencing

Mice were housed in a 12-h light/12-h dark cycle (LD) and were sacrificed six hours after light on (“DAY” group), or 18 hours after light on (“NIGHT” group).

#### Total RNA Sequencing

The construction of Illumina DNA libraries was prepared with Illumina Sample Preparation kit with rRNA depletion. Strand-specific RNAseq was done on Illumina HiSeq, 2x150 bp sequencing configuration (30 million reads per sample on average were obtained).

#### Mapping of total RNA-Seq

Paired-end reads (150bp) were aligned to the Mouse reference genome (UCSC mm10) using HISAT2 (62).

#### Quantification of RNA levels for each gene, and differential expression analysis

FeatureCounts (63) was used to quantify the number of counts for each mRNA. Differential expression analysis was performed using DESeq2 (64). low count genes (<10 reads in total) were removed before running DESeq2.

*Gene ontology analysis*. Gene ontology (GO) analysis was performed using Panther web service (65) https://pantherdb.org. Enrichment test was performed for Biological Process. The analysis performed was Statistical overrepresentation test. Categories with adjusted *P* values (Benjamini–Hochberg) smaller than 0.05 were reported.

## Data Availability

The data reported in this paper have been deposited in the Gene Expression Omnibus (GEO) database, www.ncbi.nlm.nih.gov/geo (accession no. GSE162751).

## Statistical analysis

Statistical analyses were conducted using GraphPad Prism (version 10.2.2). Statistically significant outliers were calculated using Grubb’s test for outliers. *P* ≤ 0.05 was considered statistically significant.

## Author contributions

F.M., S.G.B., and V.P. conceived and designed the project. F.S. and F.M. conceived the mouse model. P.-Y.B., A.S., F.S., J.B., M.C., D.B., F.O., C.S., F.M., A.-M.F.-B., J.K., and E.C. performed experiments. F.M., S.G.B., V.P., P.-Y.B., A.S., F.S., J.B., M.C., D.B., F.O., C.S., F.M., A.-M.F.-B., J.K., and E.C. analyzed data. P.-Y.B., A.S., A.-M..F-B., F.M., and S.G.B. wrote the manuscript. All the authors read and approved the manuscript.

## Acknowledgments

We thank Jacques Van Helden and Jean-Louis Franc for discussions. We thank the INMED and Lille animal facility platform and the INMED Molecular Biology platform. We thank Meryem Tardivel and Antonino Bongiovanni from the BICEL Photonic Microscopy in Lille. We thank members of the PLBS UAR2014-US41 for their expert technical support. SGB and FM are funded by the Foundation for Prader-Willi Research, Fédération pour la Recherche sur le Cerveau, and l’Agence National pour la Recherche (ANR-22-CE16-0007, MATBIOTA).

