## Supplementary material for "Investigation of a Novel Mouse Model of Prader-Willi Syndrome with Invalidation of *Necdin* and *Magel2*": Supplental Table and Figures

**Barelle *et al.***

**Supplemental Table and Figures**

**Supplemental Table 1. Commonly dysregulated genes in both *Madin* KO mice and PWS patients**

| DAY TF in DKO mice (27 items) | Night TF in DKO mice (69 items) | TF common in DEGs in PWS (Cell Reports) and Day TF in DKO mice (7 items) | TF common in DEGs in PWS (Cell Reports) and Night TF in DKO mice (15 items) |
|---|---|---|---|
| AHDC1 | AEBP1 | Egr3 | Aebp1 |
| CIC | ALX3 | Fosl2 | Alx3 |
| EGR3 | ARID5B | Gli1 | Arid5b |
| FOSB | ARNT | Hif3a | Etv6 |
| FOSL2 | ARNTL | Prr12 | Jund |
| FOXB1 | ARX | Rfx5 | Klf8 |
| FOXK1 | ATF1 | Sox9 | Klf9 |
| GLI1 | BAZ2B | | Npas2 |
| HES5 | BHLHE41 | | Pbx1 |
| HEYL | BNC2 | | Prox2 |
| HIF3A | EBF1 | | Rfx4 |
| HLF | ESRRA | | Scml4 |
| KMT2B | ESRRG | | Smad3 |
| MEF2D | ETV6 | | Tbx15 |
| NFIX | FLI1 | | Zbtb20 |
| NR2F6 | FLYWCH1 | | |
| OTP | FOXP1 | | |
| PRR12 | GTF3A | | |
| RAX | HDX | | |
| RFX5 | IRF2 | | |
| SALL1 | JAZF1 | | |
| SIX3 | JUND | | |
| SOX9 | KLF12 | | |
| SREBF1 | KLF8 | | |
| UNCX | KLF9 | | |
| XBP1 | LCORL | | |
| ZKSCAN2 | LIN28B | | |
| | MAF | | |
| | MBD2 | | |
| | MBD3 | | |
| | MEF2C | | |
| | NFIA | | |
| | NPAS2 | | |
| | NPAS3 | | |
| | NR1D1 | | |
| | NR3C2 | | |
| | NR6A1 | | |
| | NRF1 | | |

| | | | |
|---|---|---|---|
| | PBX1 | | |
| **DAY TF in DKO mice (27 items)** | **Night TF in DKO mice (69 items)** | **TF common in DEGs in PWS (Cell Report) and Day TF in DKO mice (27 items)** | **TF common in DEGs in PWS (Cell Report) and Night TF in DKO mice (69 items)** |
| | PRDM5 | | |
| | PRDM6 | | |
| | PRMT3 | | |
| | PROX2 | | |
| | PRRX2 | | |
| | RBPJ | | |
| | RFX4 | | |
| | RREB1 | | |
| | SCMH1 | | |
| | SCML4 | | |
| | SETBP1 | | |
| | SMAD3 | | |
| | SMYD3 | | |
| | SOX5 | | |
| | SP8 | | |
| | TBX15 | | |
| | TCF7L1 | | |
| | TEAD1 | | |
| | TERF1 | | |
| | TET2 | | |
| | THRB | | |
| | TSHZ2 | | |
| | ZBTB20 | | |
| | ZBTB7C | | |
| | ZEB1 | | |
| | ZFHX3 | | |
| | ZFPM2 | | |
| | ZMAT1 | | |
| | ZMAT4 | | |
| | ZUP1 | | |

**A** **Tests measuring Preweaning Sensorial and Motor Development**
Adapted from L. Roubertoux et al. (2018)

| | P 1 | P 2 | P 3 | P 4 | P 5 | P 6 | P 7 | P 8 | P 9 | P 10 | P 11 | P 12 | P 13 | P 14 | P 15 |
|---|---|---|---|---|---|---|---|---|---|---|---|---|---|---|---|
| **Righting response** | | | X | X | X | X | X | X | X | | | | | | |
| **Cliff avoidance** | | | | X | X | X | X | X | | | | | | | |
| **Adult paw position** | | | | | X | X | X | X | X | X | | | | | |
| **Adult walking pattern** | | | | | | X | X | X | X | X | | | | | |
| **Reaction to slope** | | | | | | X | X | X | X | X | | | | | |
| **Rooting reflex** | | | | | | | X | X | X | X | X | X | X | X | |
| **Vertical climbing** | | | | | | | | | X | X | X | X | X | X | |
| **Climb the slope** | | | | | | X | X | X | X | X | | | | | |
| **Bar holding** | | | | | | | | | | X | X | X | X | X | |
| **Eyelid opening** | | | | | | | | | | | | | | X | X |
| **Opening auditory canal** | | | | | | | | | | | | | | X | X |

**B** **Tests measuring behavior at adulthood**

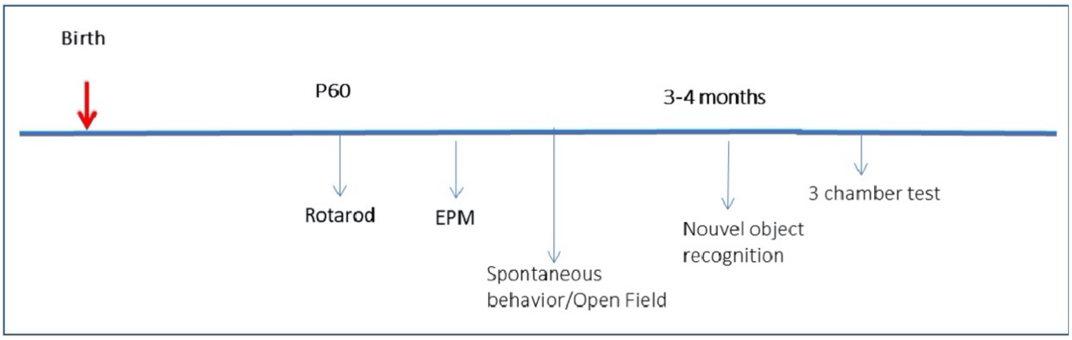


**Supplemental Figure 1. Behavioral study paradigm.** (**A**) Table listing the eleven tests performed to evaluate preweaning sensorial and motor development in *Madin* KO and WT pups between postnatal day (P) 3 and P15. (**B**) Timeline of behavioral tests perform in adult *Madin* KO and WT mice. *$P$<0.05, **$P$<0.01.

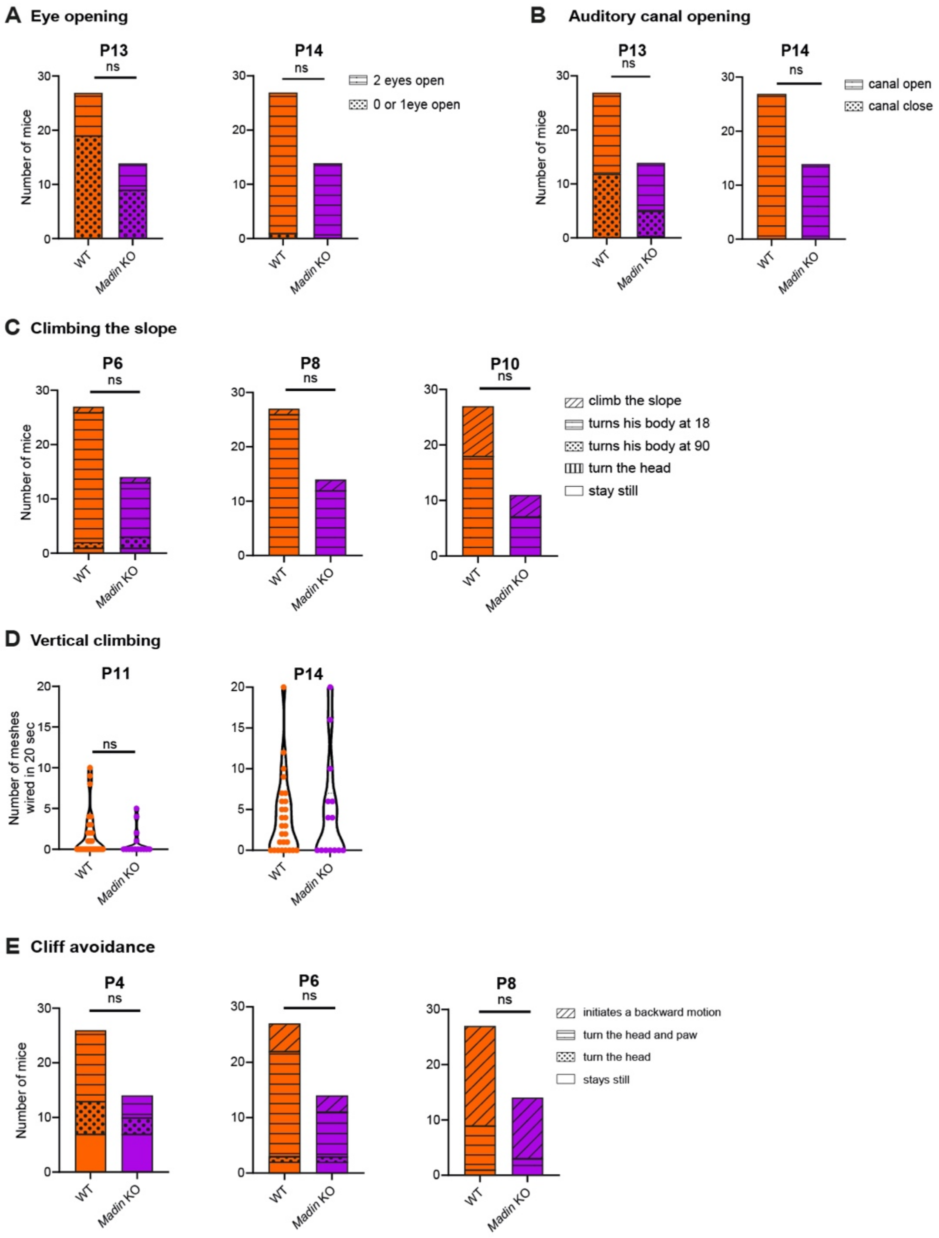


**Supplemental Figure 2. Behavioral tests performed in the two first weeks of postnatal life in *Madin* KO and WT mice.** (**A**) Number of pups displaying eye opening and (**B**) auditory canal opening at P13 and P14 (n = 14-27 animals per group). (**C**) Climbing 30° slope test in pups at P6, P8, and P10 and (**D**) vertical climbing test in pups at P11 and P14 (n = 14-27 animals per group). (**E**) Cliff avoidance test in pups at P4, P6, and P8. Statistical significance between groups was determined by a Chi$^2$ test (A,B,C,E), or a Mann-Whitney test (D).

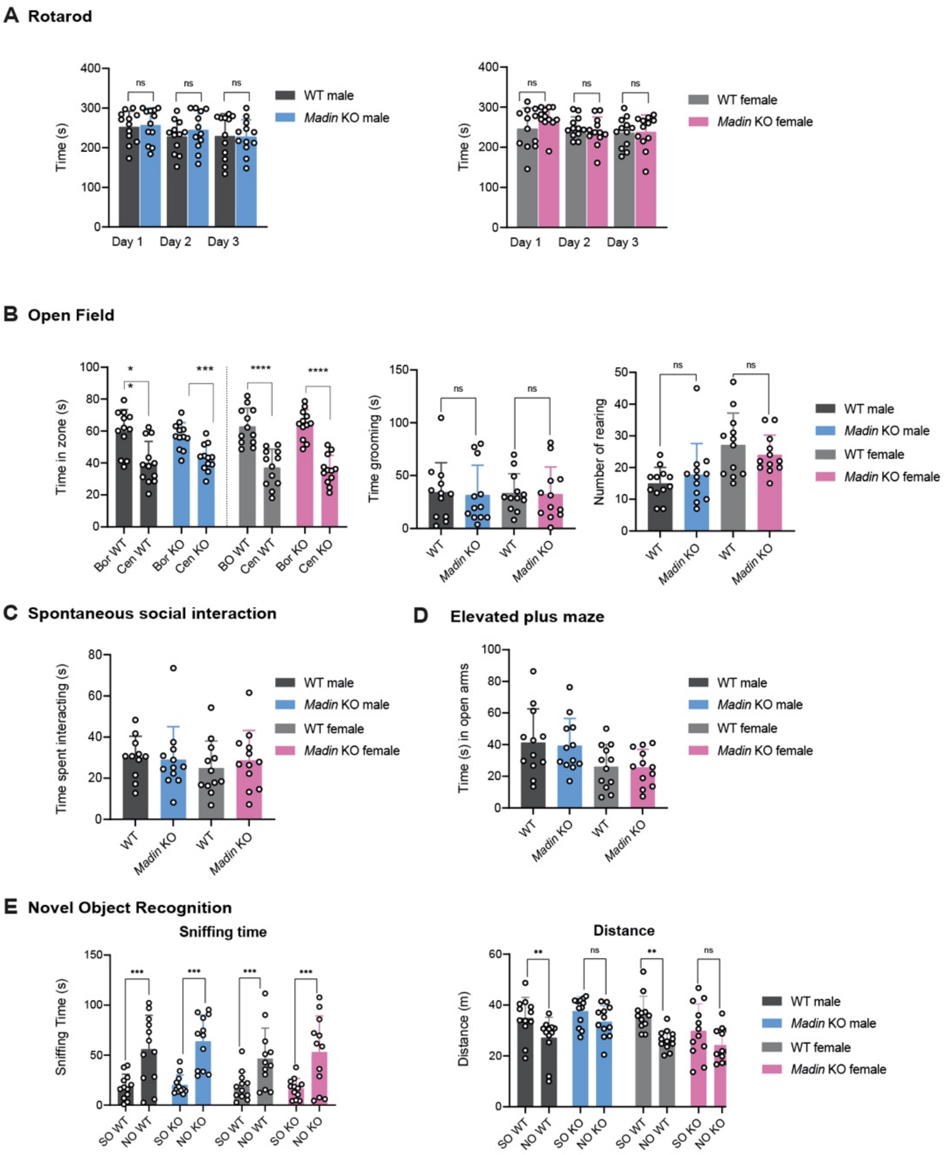


**Supplemental Figure 3. Motor ability and spontaneous, anxiety and social behaviors in adult *Madin* KO and WT mice.** (**A**) Rotarod, (**B**) open field, (**C**) spontaneous interaction, (**D**) elevated plus maze, and (**E**) novel object recognition tests in adult WT and *Madin* KO mice (n = 12 animals per group). Data are presented as mean ± SEM. Statistical significance between groups was determined by a two-way repeated measures ANOVA (A), a Mann-Whitney test (B,C,D), or a Wilcoxon matched pairs test (E). *$P < 0.05$, **$P < 0.01$.

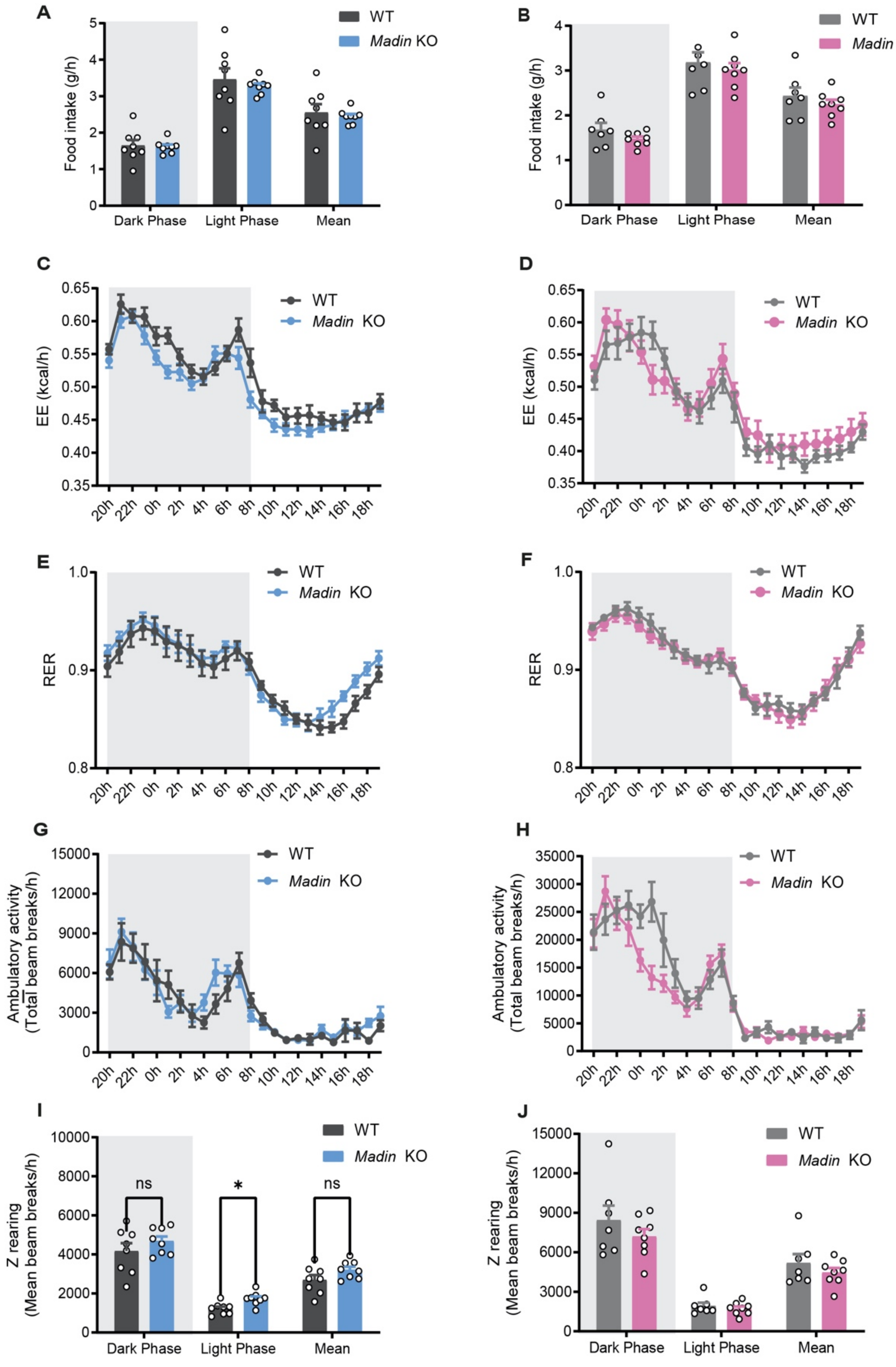


**Supplemental Figure 4. *Madin* KO mice display normal regulation of energy balance.** (**A, B**) Food intake, (**C, D**) energy expenditure, (**E, F**) respiratory exchange ratio (RER), (**G, H**) spontaneous locomotor activity (xy), and (**I, J**) number of z-rearing in (**A, C, E, G, I**) male and (**B, D, F, H, J**) female *Madin* KO and wild-type mice at P180 (n = 7-8 animals per group). Data are presented as mean ± SEM. Statistical significance between groups was determined by a Mann-Whitney test (A,B,I,J), a 2-way ANOVA followed by an uncorrected Fisher's least significant difference (LSD) test (C,D,E,F,G,H). $^{*}P \leq 0.05$.

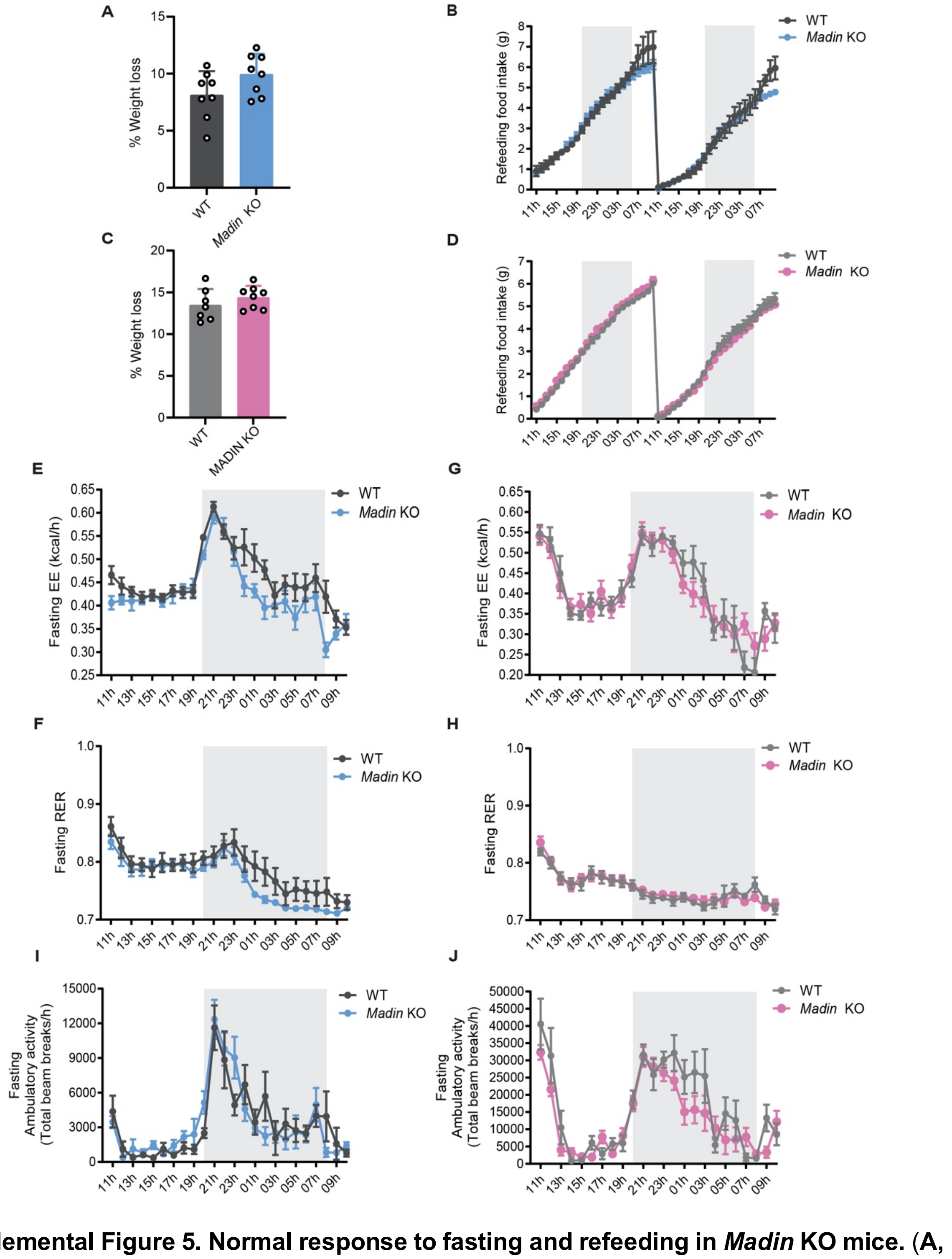


**Supplemental Figure 5. Normal response to fasting and refeeding in *Madin* KO mice.** (**A, C**) % weight loss after fasting, (**B, D**) cumulative food intake after refeeding, (**E, G**) energy expenditure during fasting, **(F, H)** respiratory exchange ratio (RER) during fasting, (**I, J**) spontaneous locomotor activity (xy), in (**A, B, E, F, I**) male and (**C, D, G, H, J**) female *Madin* KO and wild-type mice at P180 (n = 7-8 animals per group). Data are presented as mean ± SEM. Statistical significance between groups was determined by a Mann-Whitney test (A,B), a 2-way ANOVA followed by an uncorrected Fisher's least significant difference (LSD) test (B-J).
